# Atomistic molecular insight on Angiotensin-(1-7) interpeptide interactions

**DOI:** 10.1101/2023.02.19.529149

**Authors:** L. América Chi, Somayeh Asgharpour, Rodolfo Blanco-Rodríguez, Marlet Martínez-Archundia

**Affiliations:** Laboratory for the Design and Development of New Drugs and Biotechnological Innovation, Escuela Superior de Medicina, Instituto Politécnico Nacional, Plan de San Luis y Díaz Mirón, Ciudad de México 11340, Mexico; IAS-5/INM-9, Computational Biomedicine, Forschungszentrum Jülich, Wilhelm-Johnen-Strasse, 52428 Jülich; Institute for Modeling Collaboration and Innovation, University of Idaho, Moscow, Idaho, 83844–1103, USA

**Keywords:** Peptide, Ang-(1-7), Molecular Dynamics, oligomerization

## Abstract

Angiotensin-(1-7) is an endogenous peptide with vaso-protective, anti-oxidant, and anti-inflammatory effects which has been proposed as a potential therapeutic agent in a wide range of clinical conditions. Angiotensin-(1-7) presents a pH-dependent physical instability in aqueous solutions; however, it still lacks a proper atomistic study that provides insights into this behavior and its potential implications. Hence, we studied the formation of early Angiotensin-(1-7) oligomeric aggregates in an aqueous environment under acidic and neutral conditions; physiological and high ionic strength; and high and low peptide concentrations using all-atom Molecular Dynamics simulations. Our main findings are: 1) at acidic pH, there is a poor level of Angiotensin-(1-7) clustering, while, 2) at neutral pH, peptides aggregate in a unique cluster, in good trend with experimental physical instability reports and 3) an increase in salt concentration at acidic pH gives place to aggregation similar to the case at neutral pH. Our results open the route for the modulation of Angiotensin-(1-7) aggregation through a combination of salt concentration and pH conditions. Our protocol (MD + cluster analysis + amino acids interaction map analysis) is general and could be applied to other peptides to study the inter-peptide interaction mechanisms.

## 1 Introduction

Nowadays, there has been a dramatic expansion in the discovery/design of bio-active peptides^1^ and the interest in their potential use in therapeutics.^2^ Moreover, this number is likely to continue to increase in the coming years.^3,4^ However, therapeutic peptides pose considerable challenges in their aqueous formulation or their delivery in an aqueous environment, particularly concerning their physical stability.^5,6^ Physical instability implies changes to the higher-order structures, usually involving noncovalent bonds: early oligomerization, aggregation, and further precipitation. ^5^ Since peptide association to form either amorphous aggregates or highly structured fibrillar species, could limit their clinical or pharmaceutical applications, or be associated with diseases,^7^ it is crucial to study the factors affecting it.

Angiotensin-(1-7) (Ang-(1-7)) is an endogenous bio-active therapeutic heptapeptide involved in many clinical conditions ranging from cancer, cardiovascular diseases, pulmonary diseases, diabetes and COVID-19,^8,9^ which guarantees its physiological and therapeutically relevance. Ang-(1-7) shows a *β*-sheet structure in solution, with a bend stabilized by interactions between residues Val3 and Tyr4 according to circular dichroism and NMR data.^10^ Its conformational phase space has been studied theoretically, showing that the peptide is very flexible in solution.^10,11^ Although, there is not a systematic study about Ang-(1-7) solubility range in water, several reports point to solubility at least in a range from 1 *μ*M to 53 mM.^10,12–14^ Meanwhile, saturation concentration is unknown. The critical aggregation concentration for Ang-(1-7) (below which the species are predominantly in monomeric form) is thought to be around 1 mM. ^10^ Regarding its stability in an aqueous environment, a pH-dependant and time-dependent stability has been reported for Ang-(1-7) in cardio-protective aqueous solutions: at acidic and basic pH, there is no significant instability of Ang-(1-7) solutions at least until the day 25, however, at neutral pH, the solute concentration decreases to half since the first 5 days according to fluorescence experiments; once discarding chemical or bacterial degradation, Zetasizing experiments showed that the solutions at neutral pH exhibited polydispersity, indicating the formation of microparticles (regardless of the solution concentration studied). ^12^ Concerning the peptide delivery, it has been reported that in a multi-Ang-(1-7)/dendrimer system, peptide-peptide interactions collaborate with the complex formation^15^ and consequently might have implications on peptide delivery. It is clear that an in-depth understanding of the underlying mechanisms of Ang-(1-7) peptides association (early oligomerization) and the main factors modulating it are, therefore, essential to design rational strategies to optimize the stability of the peptide in an aqueous formulation or its release in an aqueous environment.

Here, we performed extensive, unbiased, explicit solvent, full-atomistic, statistical mechanics-based molecular dynamics (MD) simulations to investigate the structural and dynamical features of early oligomeric aggregates formed by Ang-(1-7) peptides. For the analysis, we determine the oligomer size and inter-peptide contacts that drive the early aggregation process. As a peptide in an aqueous solution can be stabilized against aggregation by optimizing the pH and ionic strength of the solution, acidic and neutral pH conditions, and physiological and high salt concentrations conditions were tested. To the best of our knowledge, this is the first atomistic study on Ang-(1-7) peptide-peptide interactions in an aqueous solution performed so far. Our results correlated well with experimental instability tendencies and gave insight into structural and dynamical features of oligomeric early aggregates at atomistic resolution.

## 2 Methods

### Molecular model systems

The solution-NMR structure of Ang-(1-7) peptide (ASP-ARG-VAL-TYR-ILE-HIS-PRO) was retrieved from the protein database (PDB) with ID: 2JP8; the topology was created using the pdb2gmx module of GROMACS-2019.2+plumed-2.6.0^16–19^ and their parameters were downloaded from the GROMOS-compatible 2016H66 force field^20^ or CHARMM27 force field;^21^ protonation states were assigned according to physical-chemical properties predicted with peptide calculator server^22^ and verified by using ProPkA software^23^ (zero peptide charge for neutral pH and +2 for acidic conditions) as reported in our previous work. ^11^

### MD of peptides in solution

To consider the possible influence of the conformation and placement (position + orientation) of the initial peptides on the aggregation states, several initial peptide conformations were used and all the important simulations were performed in duplicate starting from different initial peptides placements. The different initial peptide conformations were taken from the lowest energy models reported at the NMR experiments. The necessary number of peptide conformers were randomly placed on a cubic simulation box. Then the system was solvated by adding water molecules. Ions were added for neutralizing the net charge of the system and mimicking two different ionic concentrations, 0.15 M and 1 M (named as low and high ionic concentrations, respectively). The box dimensions were chosen to give two peptide concentrations of about 12 mM and 34 mM (named as low and high solute concentrations, respectively). This decision was made considering concentrations above the suggested critical aggregation concentration, ^10^ above the concentrations tested in instability studies, within the solubility concentrations reported, ^12^ as well as concentrations that allow a reasonable system size according to the computational resources available. Finally, the systems were subjected to minimization, equilibration and production of standard MD simulations using GROMACS-2019.2+plumed-2.6.0^16–19^ software. Details on the molecular systems considered in the present work are presented in Table 1.

**Table 1:** Molecular systems considered in the present work. “A” means Angiotensin-(1-7) peptides, the first super index could be acidic (a) or neutral (n) conditions, the second super index could be low (l) or high (h) solute concentration, the third super index could be low (l) or high (h) salt concentration and the last super index is related with CHARMM27 (c) or GROMOS-based 2016H66 (g) force fields used. Simulations of systems performed at 34 mM and with CHARMM27 force field were performed in duplicate.

| Type | pH | Concentration<br>[mM] | Peptides<br>[No.] | Ions<br>[M] | Cl<sup>-</sup><br>[No.] | Na<sup>+</sup><br>[No.] | Waters<br>[molecules] | Total<br>[atoms] | Box-side<br>[nm] |
| --- | --- | --- | --- | --- | --- | --- | --- | --- | --- |
| A<sup>a,l,l,c</sup> | 3 | 12 | 6 | 0.15 | 77 | 89 | 26839 | 81448 | 9.5 |
| A<sup>a,h,l,c</sup> | 3 | 34 | 6 | 0.15 | 25 | 37 | 8604 | 26642 | 6.5 |
| A<sup>a,h,l,g</sup> | 3 | 34 | 6 | 0.15 | 25 | 37 | 8628 | 26468 | 6.5 |
| A<sup>a,h,h,c</sup> | 3 | 34 | 6 | 1 | 165 | 177 | 8324 | 26082 | 6.5 |
| A<sup>n,h,l,c</sup> | 7 | 34 | 6 | 0.15 | 25 | 25 | 8617 | 26657 | 6.5 |

### Systems nomenclature

In order to make it clear the system nomenclature is described in detail as follows:

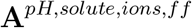

where the first superindex position is related to pH and can take values of acidic (a) or neutral (n), the second super index could be low (l) [12 mM] or high (h) [34 mM] solute (Ang-(1-7)) concentration, the third super index could be low (l) [0.15 M] or high (h) [1 M] salt concentration and the last super index is related with CHARMM27 (c) or GROMOS-based 2016H66 (g) force fields used.

### Clusters analysis

The peptide’s tendency to interact and form aggregates can be quantitatively characterized by analyzing the formation of clusters (or aggregates) during the MD trajectories. These cluster analyses were performed by using our in-house Python code, which was adapted from the work of Samantray et al. ^24^ In this code, the positions of the atoms in the peptides are read from the trajectory file obtained from MD using the MDAnalysis python library. The inter-peptide distance is taken as the shortest distance among all the distances between the atoms of one peptide with those of the other. We considered a cluster when the inter-peptide distance is less than 3 Å, and the cluster size (CS) is defined as the number of peptides forming a cluster. We analyzed the formation of clusters during the MD trajectory using the `igraph` python library; in the code, each peptide is considered as a node in a graph and when the inter-peptide distance is less than the threshold, a link is defined between the two nodes involved. Thus, for each frame of the trajectory, we have a graph where its unconnected components are clusters with the number of nodes as the cluster size; the extreme cases are that all nodes are disconnected (clusters formed by only one peptide) and a complete graph (a single cluster formed by all peptides). As we mentioned above, the cluster size could be related to the instability (or aggregation propensity) of the peptides. On the other hand, we would give molecular insights into the inter-peptide interactions if we consider residue–residue contacts between the peptides forming the clusters, hence we measure the frequency at which the amino acids between different peptides are close enough to each other, where the contact is counted if any of the atoms between two amino acids is closer than 3Å. The result of this count is reported as a probability map. An alternative for this analysis is the gromacs command `gmx mdmat`, with which we can obtain the map of contact frequencies between residues for peptides pairs, then at the end the matrices of all peptide pairs are averaged. We decided to calculate it because our code that records the clusters can give us the same information for all peptides at the same time. Our code is available on GitHub using the next link: https://github.com/rblanco89/oligomerization.git.

## 3 Computational details

During the MD simulations, each system was placed in a cubic simulation box such that all peptide atoms were, at least, 1 nm distant from the box edges. The box was then filled with a sufficiently large number of SPC water molecules. The MD simulation box was neutralized by adding the appropriate number of Cl^-^ and Na^+^ counterions at a physiological and high salt concentration, 0.15 M and 1 M, respectively. Systems were subsequently energy minimized under periodic boundary conditions. The equilibration of each system was first performed under the NVT canonical ensemble for 1000 ps. Initial velocities were generated according to a Maxwell-Boltzmann distribution corresponding to a temperature of 300 K. Pressure coupling was then switched on, and the system was equilibrated under the NPT isothermal–isobaric ensemble for 1000 ps with a reference pressure of 1 bar. Production runs were carried out under NPT ensemble at 298.15 K and 1 bar for 400 ns. The coordinates were written to the output file every 10 ps for the final analysis. Newtonian equations of motion were integrated using the leapfrog scheme with a time step of 2 fs. Periodic boundary conditions were adopted in all cases. All bond lengths were constrained to their reference values employing the LINCS algorithm. ^25^ Electrostatic interactions were calculated with the PME method.^26^ Atomic-based pair list generation was employed, as implemented in the Verlet algorithm. When needed, V-rescale^27^ thermostat at 298 K and Parrinello-Rahman^28^ barostat at 1 bar were used.

## 4 Results and analysis

Short peptides like Ang-(1-7) in solution could be hard to characterize experimentally, mainly because of their dynamical behavior when not in complex with a macromolecule. Computational methods like MD simulations offer an alternative to studying them at atomic resolution, providing insights into the underlying interactions between peptide-peptide, peptide-solvent, or peptide-ions elements in the system. Here, peptide-peptide interactions were studied using MD simulations in presence of an explicit solvent. We evaluated the effect of the pH, peptide concentration, salt concentration, and force field used on peptide aggregation propensity according to the following metrics: the radius of gyration of all peptides (R_*g*_), the number of monomers, the maximum cluster size, and peptide-peptide interaction map.

### 4.1 Radius of gyration of all peptides

The radius of gyration (R_*g*_) of all the peptides present in an aqueous system gives us a measure of the compactness of the peptide arrangement, which can be related to the degree of peptides’ clustering (aggregation) as demonstrated in previous works. ^29^ Importantly, the calculation of R_*g*_ of a system of peptides requires knowledge of the center of mass of the complete system. However, in a simulation box with periodic boundary conditions, the center of mass of the peptide system depends on which periodic image of the peptides near the boundary we take so that we may have different R_*g*_ according to the chosen image. Because of this drawback, we considered calculating the R_*g*_ of the peptide system, placing the reference point on the center of mass of each peptide, and averaging the results. Therefore, the time evolution of the mean R_*g*_ for each system tested here is presented in Figure 1.

**Figure 1:**
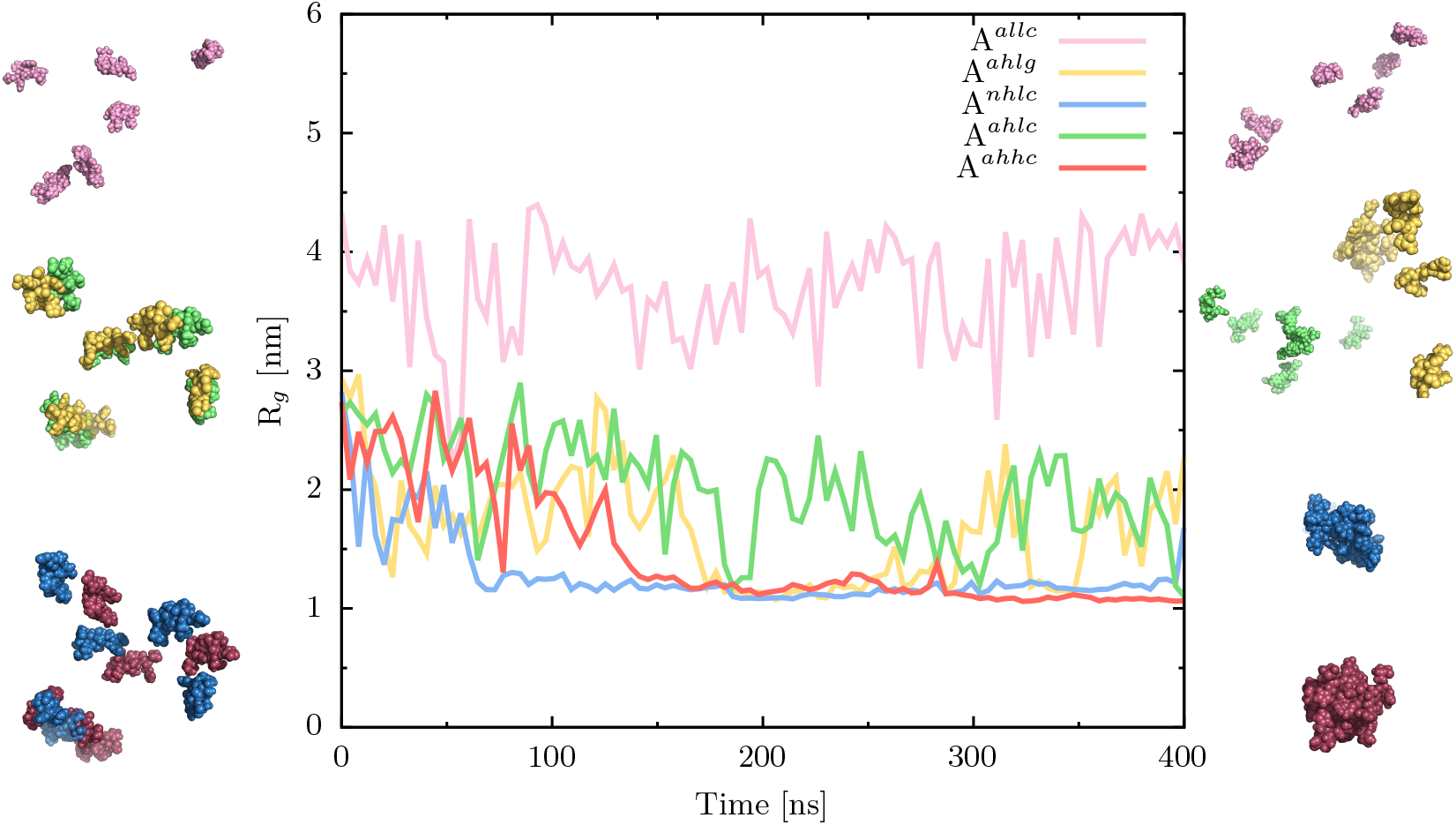
The Radius of gyration (R_*g*_) of all peptides as a function of time. Each line shows the average values of the R_*g*_ when the reference center of mass is considered on a different peptide. A splines-based smooth fitting was applied for the sake of visibility. Initial frames (left) and representative frames of the equilibrated region (right) are presented. “A” means Angiotensin-(1-7) peptides, the first super index indicates either acidic (a) or neutral (n) conditions, the second super index indicates either low (l) or high (h) solute concentration, the third super index indicates either low (l) or high (h) salt concentration, and the last super index is related to CHARMM27 (c) or GROMOS-based 2016H66 (g) force fields used.

We ran simulations at two different Ang-(1-7) concentrations (12 mM and 34 mM), leaving all the other variables constant (acidic pH, 15 mM NaCl and CHARMM ff), these systems are represented as A^*a,l,l,c*^ and A^*a,h,l,c*^, respectively. What we can observe from their R_*g*_ comparison is that they both show different levels of compactness, do not form a big unique cluster (R_*g*_ between 3 and 4.5 nm in case of A^*a,l,l,c*^; R_*g*_ between 1 and 3 nm in case of A^*a,h,l,c*^) during most parts of the simulation (they do not tend to aggregate), which could be associated with different levels of clustering (monomers, dimers, trimers, etc). This might be related to the electrostatic repulsion between the positively charged peptides at this pH (each peptide has a charge of +2).

We ran simulations at two different force fields (CHARMM27 and GROMOS-based 2016H66 ff), leaving all the other variables constant (acidic pH, 34 mM Ang-(1-7) and 15 mM NaCl), these systems are represented as A^*a,h,l,c*^ and A^*a,h,l,g*^, respectively. GROMOS-based 2016H66 and CHARMM27 force fields were previously used to simulate Ang-(1-7),^11,15^ additionally CHARMM27 is a popular and well-established force field to simulate amino-acid-based molecules and short peptides.^30^ In the case of the GROMOS-based 2016H66 ff, we observed a significant increment in the unique cluster state (R_*g*_ around 1 nm), but still, it went back and forth between different cluster levels during the simulation time as for A^*a,h,l,c*^ and A^*a,l,l,c*^ systems. It might mean that GROMOS-based 2016H66 ff could slightly overestimate the aggregation propensity of Ang-(1-7), but further investigation is needed in this regard.

We ran simulations at two different pH conditions (neutral and acidic), leaving all the other variables constant (34 mM Ang-(1-7), 15 mM NaCl and CHARMM ff), these systems are represented as A^*n,h,l,c*^ and A^*a,h,l,c*^, respectively. We observed that at neutral pH, there is a predominant R_*g*_ state since the first 100 ns (R_*g*_ around 1 nm), presumably belonging to the one unique cluster state organic-rich phase. This behavior seems to be in line with the instability reported experimentally, showing a marked propensity to aggregate at neutral pH but not at acidic or basic conditions. ^12^ This might make sense due to the neutrally charged peptides (isoelectric point = 7.8) at this pH compared with basic or acidic conditions. As the critical concentration of Ang-(1-7) is thought to be about 1 mM at neutral pH, it was expected that all peptides in the simulated system belong to the organic-rich phase.

Finally, we ran simulations at two different NaCl salt concentrations (physiological and salty conditions), leaving all the other variables constant (acidic, 34 mM Ang-(1-7), and CHARMM ff), these systems are represented as A^*n,h,l,c*^ and A^*a,h,h,c*^, respectively. We observed that the behavior of the salty system is very similar to the one at neutral pH (R_*g*_ around 1 nm) from the first 150 ns. This could be related to the Hofmeister^31^ effect that takes place at higher salt concentrations, where kosmotropes (like Na^+^ and to less extent Cl^-^) enhance the hydrophobic effect of the solute by tightening the structure of water around the ions, leading to an increase in VdW interactions between peptides or the well-known effect triggered by an increase in the salt concentration of a solution which results in the shielding off of the protein’s charge leading to a reduction in the electrostatic repulsion of the proteins among themselves, which results in their aggregation and precipitation. ^32^ At low salt concentrations (~ 10 mM), the Debye–Huckel theory explains that solute molecules are surrounded by the salt counter ions, and this screening results in decreasing electrostatic free energy between solute molecules and increasing activity of the solvent-solute interactions, which in turn leads to increased solubility. These results could open the possibility to modulate Ang-(1-7) peptide-peptide interactions through salt concentration.

From the five systems tested, two of them seem to be prone to aggregation: systems under neutral pH and under high salt concentration, A^*n,h,l,c*^ and A^*n,h,h,c*^, respectively. The rest of the systems show a differential behavior that shows less aggregation tendency. In previous works, it was shown that a full atom molecular dynamics simulation of a multiple peptide system can be in line with experimentally determined solubility values^33^ or peptide aggregation propensity values^29^ and here our results seem to confirm this potential.

### 4.2 Key players on aggregation

Ions play a key role in Ang-(1-7) behavior in water. As can be seen in Figure 2a, the radial distribution function (RDF) between Cl^-^ and peptides is differential according to pH and ionic salt concentration. In the case of A^*a,h,l,c*^ system (acidic pH conditions, and physiological salt concentration) we observed a bigger probability for the formation of ionic bonds between the Cl^-^ anions and the positively charged moieties on peptides (N-terminal, ARG^+^ and HIS^+^) at the first three well-defined peaks around 0.2, 0.3 and 0.4 nm. In the case of A^*n,h,l,c*^ system (neutral pH conditions, and physiological salt concentration) we observed less probability for the formation of ionic bonds between the Cl^-^ anions and the positively charged moieties on peptides. Additionally, the RDF peak at 4 nm almost disappears, this peak is related to the HIS residue, showing a crucial role of HIS protonation on Ang-(1-7) aggregation. The other significant change in Ang-(1-7) peptides at different pH conditions are the C-terminal neutralization at acidic pH compared with the negative charge at neutral pH. It appears that the Ang-(1-7) C^-^-terminus is not only crucial for Ang-(1-7) complexation with its MAS receptor in cells through electrostatic interaction with ARG245 on MAS^34^ but could also have implications on the modulation of aggregates formation. In the case of A^*a,h,h,c*^ system (acidic pH conditions, and high salt concentration) we observed less probability for the formation of ionic bonds between the Cl^-^ anions and the positively charged moieties on peptides as in the case of neutral pH. In general, neutral and salty systems show similar behavior when interacting with Cl^-^ neighbors, and this could be related to their aggregation in a unique cluster compared with acidic conditions where there is not a marked organic-rich phase. As can be seen in Figure 2b, the radial distribution function (RDF) between Cl^-^ and Na^+^ is differential according to pH and ionic salt concentration. In the case of A^*n,h,l,c*^ system (neutral pH conditions, and physiological salt concentration) we observed a bigger probability for the formation of ionic bonds between the Cl^-^ anions and Na^+^ cations in comparison with acidic or salty systems. That means that even though Cl^-^ anions are less dense around peptides at neutral and salty conditions, they play different roles in water, preferring to form ion pairs with Na^+^ cations in the case of neutral pH but staying surrounded by waters in salty systems.

**Figure 2:**
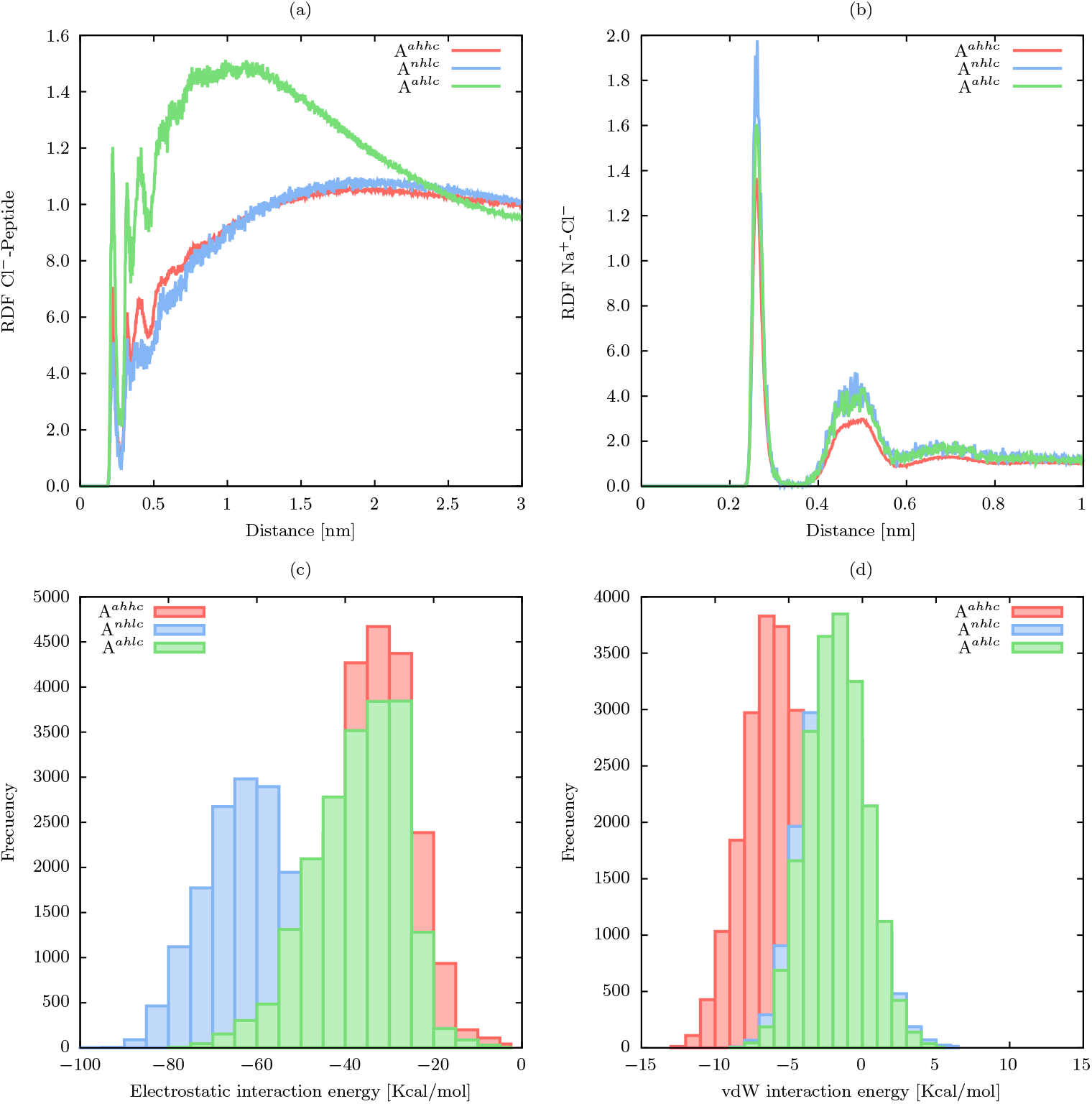
RDF between Cl^-^ and peptides or Na^+^; and energy contributions predominated in the Ang-(1–7)/Ang-(1-7) interaction. (a) RDF between Cl^-^ and peptides,(b) RDF between Cl^-^ and Na^+^ (c) frequency plot of the electrostatic interaction energy between Ang-(1–7)/Ang-(1–7), in the final 200 ns of MD simulation and (d) frequency plot of the vdW interaction energy between Ang-(1–7)/Ang-(1–7), in the final 200 ns of MD simulation. Interaction energy represents an average value for the 6 peptides. Abbreviations: “A” means Angiotensin-(1-7) peptides, the first super index indicates either acidic (a) or neutral (n) conditions, the second super index indicates either low (l) or high (h) solute concentration, the third super index indicates either low (l) or high (h) salt concentration, and the last super index is related to CHARMM27 (c) or GROMOS-based 2016H66 (g) force fields used.

To have insight into the driving forces governing the interactions between peptides, the electrostatic and vdW interaction energies between Ang-(1–7)-Ang-(1–7) peptides were calculated during the last 200 ns of MD simulations. Regarding the nature of the interactions driving aggregation, electrostatic energy played a major contribution as can be seen in Figure 2c-d for all three cases: acidic, neutral, and salty systems. This result suggests that the interactions of charged and polar amino acids could be the primary forces driving oligomerization. In the case of vdW interactions, we observed that they are also important in inter-peptide interactions though to a less extent compared with electrostatic interactions, especially due to hydrophobic residues (ILE, PRO, VAL, TYR). These results are supported by a previous study^15^ where they found that peptide/peptide interactions displayed high electrostatic interaction energy values, and collaborated in complex formation with a dendrimer by performing MD simulations on neutral pH conditions and CHARMM27 ff. In salty system, the vdW interactions become more relevant, supporting that according to the Hofmeister^31^ effect, at higher salt concentrations, kosmotropes (like Na^+^ and to less extent Cl^-^) enhance the hydrophobic effect of the solute by tightening the structure of water around the ions, leading to an increase in VdW interactions between peptides. This could be the reason why at high salt concentrations, Ang-(1-7) peptides aggregate.

### 4.3 Cluster analysis

To clarify further the R_*g*_ results, the cluster analysis addresses two kinds of information: 1) the first one is related to the quantity of smaller entities (monomers) and the size of the biggest (maximum cluster size) entities present at each frame, and 2) the distribution of different cluster sizes during the simulation in the equilibrated region. Therefore, the time evolution of the number of monomers, the time evolution of the maximum cluster size, and the cluster size distributions for the whole simulation are displayed in Figure 3 for each one of the systems tested in this work. The number of monomers and the maximum cluster size in the last 200 ns of the MD simulation are displayed in Figure 3a and 3b, respectively. In these plots, the signals were filtered using convolution with a step function with a window of 25 ns. Figure 3a shows that the number of monomers of A^*a,l,l,c*^ ranges around 4, the highest number compared with the other systems. For A^*a,h,l,g*^ and A^*a,h,l,c*^ systems, the number of monomers ranges between 0 and 3, while the other systems show that the number of monomers is zero. Figure 3b shows that the maximum cluster size for A^*a,l,l,c*^ ranges around 2; for A^*a,h,l,g*^ and A^*a,h,l,c*^ systems it ranges between 2 and 6; and for A^*n,h,l,c*^ and A^*a,h,h,c*^ systems, the largest cluster is 6 most of the time.

**Figure 3:**
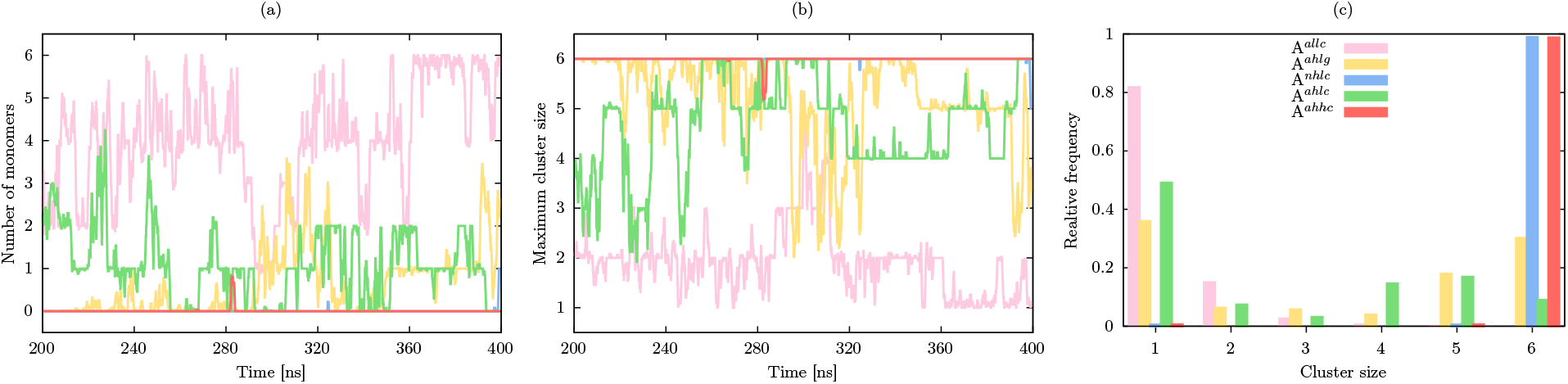
(a) Number of monomers, (b) maximum cluster size, and (c) cluster size distributions for the systems A^*a,l,l,c*^, A^*a,h,l,g*^, A^*n,h,l,c*^, A^*a,h,l,c*^, and A^*a,l,h,c*^. “A” means Angiotensin-(1-7) peptides, the first super index indicates either acidic (a) or neutral (n) conditions, the second super index indicates either low (l) or high (h) solute concentration, the third super index indicates either low (l) or high (h) salt concentration, and the last super index is related with CHARMM27 (c) or GROMOS-based 2016H66 (g) force fields used.

Figure 3c shows the distribution of clusters throughout the last 200 ns of the MD simulation, the maximum size can reach 6 (all peptides together) and the minimum is 1 (a single peptide). A^*a,l,l,c*^ system shows that during the whole simulation, the cluster size is inversely proportional to the relative frequency of occurrence. Finding a dimer and 4 monomers is a highly probable state for this system. A^*a,h,l,g*^ system shows that its cluster size is distributed mainly at the monomeric state, followed by the hexameric state and pentameric state. Finding the one-monomer/one-pentamer state, and the one-hexamer state are almost equally probable for this system. Still, the different intermediate configurations have a considerable probability as well. A^*n,h,l,c*^ system shows that the more favorable cluster size is 6. Confirming that above the critical aggregation concentration and neutral pH, all peptides in the simulated system belong to the organic-rich phase. Meanwhile, the water-rich phase of such a solution is composed of pure water. This could also be related to the macroscopic instability found for Ang-(1-7) at neutral pH. A^*a,h,l,c*^ system shows that its cluster size is distributed mainly at the monomeric state followed by the pentameric, the tetrameric, and the hexameric states. Finding a one-monomer/one-pentamer state is highly probable for this system as well as the different intermediate configurations. This is a similar behavior as the one observed for the A^*a,h,l,g*^ system, but the one hexamer state alone is much less favorable, with similar relativity frequencies as pentameric and tetrameric states. What is clear is that in those systems, peptides are not as prone to aggregate in a unique cluster as in the case of neutral pH. This could also be related to the macroscopic stability found for Ang-(1-7) at acidic and basic pH. A^*a,h,h,c*^, system shows that the more favorable cluster size is 6. Mimicking the behavior observed at neutral pH but with low salt concentration, all peptides in the simulated system belong to the organic-rich phase, meanwhile, the water-rich phase of such a solution is composed of pure water. In summary, our results make sense qualitatively with the pH-dependant experimental instability reported for Ang-(1-7) in aqueous solutions: we can show here that even in the case where there is instability (acidic pH), peptides tend to interact but in less degree, forming smaller clusters compared to that in which they form part of the organic-rich phase (neutral pH). Additionally, these results open the possibility to modulate Ang-(1-7) aggregation through salt concentration in cases where Ang-(1-7) stability is required like in the case of cardioprotective solutions.^12^ It also opens the possibility to modulate Ang-(1-7) inter-peptide interaction contribution to the load/unload into macromolecules through salt concentration.^11,15^ Ang-(1-7) aggregation does not have a documented implication in disease but it is clear that peptides are prone to interact to a different degree under the conditions tested here and could be related to the detrimental effect of Ang-(1-7) peptide at high concentrations where overdosing might interfere with its receptor-associated functions. ^35^

### 4.4 Contact analysis

The residue–residue contacts between the peptides composing the oligomers are counted and reported as a probability map in Figure 4.

**Figure 4:**
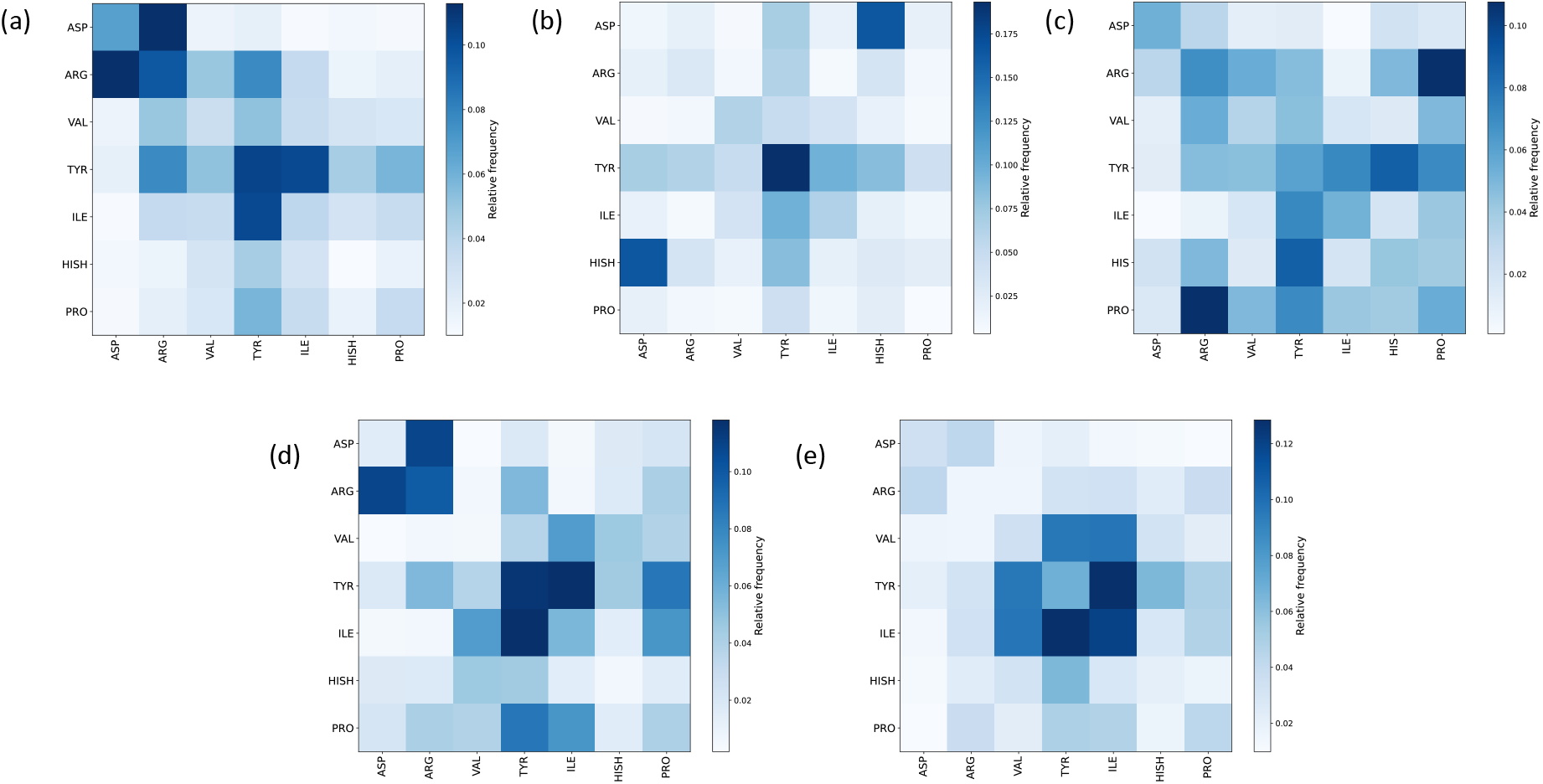
The inter-residue contact map with probabilities according to the color scale on the right for (a) A^*a,l,l,c*^, (b) A^*a,h,l,g*^, (c) A^*n,h,l,c*^, (d) A^*a,h,l,c*^, and (e) A^*a,h,h,c*^ systems. Where, “A” means Angiotensin-(1-7) peptides, the first super index could be acidic (a) or neutral (n) conditions, the second super index could be low (l) or high (h) solute concentration, the third super index could be low (l) or high (h) salt concentration and the last super index is related with CHARMM27 (c) or GROMOS-based 2016H66 (g) force fields used.

Figure 4a, A^*a,l,l,c*^ system, shows that peptides make a higher number of contacts at ASPASP, ARG-ASP, TYR-TYR, and ILE-TYR residues, which might be related to strong interactions at these positions. This residue-residue contact arrangement seems to imply a quasi-parallel orientation stabilized by electrostatic interactions between the positively charged ASP N-terminal of one peptide and the negatively charged side chain of ASP residue of another peptide; the positively charged side chain of ARG on one peptide, and the negatively charged ASP N-terminal of the other peptide; and strong hydrophobic contacts between TYR-TYR and TYR-ILE side chains.

Figure 4b, A^*a,h,l,g*^ system, shows that peptides make a higher number of contacts at HISH-ASP, TYR-TYR, and ILE-TYR, ILE-VAL, and ILE-ILE residues, which might be related to strong interactions at these positions. This residue-residue contact arrangement seems to imply a quasi-anti-parallel orientation stabilized by electrostatic interactions between the positively charged HISH side chain of one peptide and the negatively charged side chain of ASP of another one, and strong hydrophobic contacts between TYR-TYR, and ILE-TYR, ILE-VAL, ILE-ILE side chains.

Figure 4c, A^*n,h,l,c*^ system, shows that peptides make higher number of contacts at PROARG, ASP-ASP, PRO-TYR, HIS-TYR, TYR-TYR, and ILE-TYR residues, which might be related to strong interactions at these positions. This residue-residue contact arrangement seems to imply a quasi-parallel orientation stabilized by electrostatic interactions between the negatively charged ASP N-terminal of one peptide and the negatively charged ASP side chain of another peptide, the negatively charged C-terminal of PRO residue on one peptide and, the positively charged ARG side chain of another peptide, the positively charged side chain of ARG in one peptide and the negatively charged N-terminal of ASP on another peptide, strong hydrophobic contacts are formed between PRO-TYR, HIS-TYR, TYR-TYR, and ILE-TYR side chains.

Figure 4d, A^*a,h,l,c*^ system, shows that peptides make a higher number of contacts at ARG-ASP, ARG-ARG, TYR-TYR, PRO-TYR, and ILE-TYR residues, which might be related to strong interactions at these positions. This residue-residue contact arrangement seems to imply a quasi-parallel orientation stabilized by electrostatic interactions between the positively charged ARG side chain of one peptide and the negatively charged ASP N-terminal of the other peptide, and strong hydrophobic contacts between TYR-TYR, TYR-ILE, and PRO-TYR side chains.

Figure 4e, A^*a,h,h,c*^ system, shows that peptides make a higher number of contacts at HIS-TYR, TYR-VAL, and ILE-TYR, ILE-VAL, and ILE-ILE residues, which might be related to strong interactions at these positions. This residue-residue contact arrangement seems to imply a quasi-anti-parallel orientation stabilized by strong hydrophobic contacts formed between HIS-TYR, TYR-VAL, TYR-TYR, and ILE-TYR, ILE-VAL, ILE-ILE residues of different peptides.

## 5 Conclusions

The main behavior observed is that Ang-(1-7) peptides tend to aggregate more at neutral pH than at acidic pH in trend with the instability observed experimentally neat the neutral pH conditions.

We provide insight into how Ang-(1-7) peptides interact within themselves under different conditions. The more frequent residue-residue contacts occur between TYR-TYR and ILE-TYR residues in all systems. All systems appear to have electrostatic contributions as well as hydrophobic contributions to the peptide-peptide interactions. Our results validate the high electrostatic interaction energy values of peptide-peptide interactions displayed at neutral pH which were reported previously in the computational work.

We also evaluated the effect of salt concentration increment on these Ang-(1-7) peptide-peptide interactions, showing that ionic strength plays a key role in Ang-(1-7) aggregation propensity and opening the possibility to modulate its solubility/aggregation changing salt concentration in the solution.

Additionally, one of the main concerns when performing MD simulations of molecules is whether the force field and the simulation protocol can correctly model the main properties we are interested in. In the case of multi-peptide loaded intro macromolecule systems like the one presented in our previous work, where 3 Ang-(1-7) peptides were loaded into PAMAM dendrimers, we aimed at testing not only if the force field and protocol were able to reproduce structural features of the isolated peptide and PAMAM dendrimers, but also if reasonable peptide-peptide interactions were reproduced. In this work, we observed that GROMOS-based 2016H66 and CHARMM27 have a small discrepancy when forming clusters at acidic pH, either GROMOS-based 2016H66 is overestimating aggregation propensity or CHARMM27 is underestimating it, but due to the fact that the results from CHARMM27 ff appear to be more in line with the instability observed in experimental results and that it is a very used and popular force field for amino-acid molecules simulations, we think it could better model the Ang-(1-7) inter-peptide interactions.

If the data about solubility/aggregation/instability is available, it could become a standard protocol to study inter-peptide interactions to validate its modeling for aggregation-prone peptide diseases, coupling of therapeutic multiple peptides into macromolecules or in this case getting an atomic molecular insight of peptide-peptide interactions. Here, we provide a protocol along with freely available python scripts, easy to adapt to new systems that have not been provided before. The interactions’ contact map script is even useful in cases where we want to analyze interactions between identical proteins or homodimers.

## References

(1) Daliri, E. B.-M.; Lee, B. H.; Oh, D. H. Current trends and perspectives of bioactive peptides. Critical Reviews in Food Science and Nutrition 2018, 58, 2273–2284.

(2) Fosgerau, K.; Hoffmann, T. Peptide therapeutics: current status and future directions. Drug Discovery Today 2015, 20, 122–128.

(3) Wang, L.; Wang, N.; Zhang, W.; Cheng, X.; Yan, Z.; Shao, G.; Wang, X.; Wang, R.; Fu, C. Therapeutic peptides: current applications and future directions. Signal Transduction and Targeted Therapy 2022, 7, 1–27.

(4) Lau, J. L.; Dunn, M. K. Therapeutic peptides: Historical perspectives, current development trends, and future directions. Bioorganic & Medicinal Chemistry 2018, 26, 2700–2707.

(5) Avanti, C. Innovative strategies for stabilization of therapeutic peptides in aqueous formulations; 2012.

(6) Zapadka, K. L.; Becher, F. J.; Gomes dos Santos, A.; Jackson, S. E. Factors affecting the physical stability (aggregation) of peptide therapeutics. Interface Focus 2017, 7, 20170030.

(7) Armiento, V.; Spanopoulou, A.; Kapurniotu, A. Peptide-Based Molecular Strategies To Interfere with Protein Misfolding, Aggregation, and Cell Degeneration. Angewandte Chemie International Edition 2020, 59, 3372–3384.

(8) Iusuf, D.; Henning, R. H.; van Gilst, W. H.; Roks, A. J. Angiotensin-(1–7): pharmacological properties and pharmacotherapeutic perspectives. European Journal of Pharmacology 2008, 585, 303–312.

(9) Khajehpour, S.; Aghazadeh-Habashi, A. Targeting the protective arm of the reninangiotensin system: Focused on angiotensin-(1–7). Journal of Pharmacology and Experimental Therapeutics 2021, 377, 64–74.

(10) Lula, I.; Denadai, Â. L.; Resende, J. M.; de Sousa, F. B.; de Lima, G. F.; Pilo-Veloso, D.; Heine, T.; Duarte, H. A.; Santos, R. A.; Sinisterra, R. D. Study of angiotensin-(1– 7) vasoactive peptide and its *β*-cyclodextrin inclusion complexes: complete sequencespecific NMR assignments and structural studies. Peptides 2007, 28, 2199–2210.

(11) Chi, L.; Asgharpour, S.; Correa-Basurto, J.; Bandala, C. R.; Martínez-Archundia, M. Unveiling the G4-PAMAM capacity to bind and protect Ang-(1-7) bioactive peptide by molecular dynamics simulations. Journal of Computer-aided Molecular Design 2022, 1–23.

(12) Russ, M.; Jauk, S.; Wintersteiger, R.; Gesslbauer, B.; Greilberger, J.; Andrä, M.; Ortner, A. Stabilization of Angiotensin-(1-7) in Cardioprotective Solutions. International Journal of Peptide Research and Therapeutics 2019, 25, 1271–1278.

(13) KGaA, M. Angiotensin Fragment 1-7 acetate salt hydrate. 2022; https://www.sigmaaldrich.com/MX/es/product/sigma/a9202.

(14) Technology, A. Angiotensin (1-7). 2013; https://www.apexbt.com/angiotensin-1-7.html#cite.

(15) Márquez-Miranda, V.; Abrigo, J.; Rivera, J. C.; Araya-Duran, I.; Aravena, J.; Simon, F.; Pacheco, N.; González-Nilo, F. D.; Cabello-Verrugio, C. The complex of PAMAM-OH dendrimer with Angiotensin (1–7) prevented the disuse-induced skeletal muscle atrophy in mice. International Journal of Nanomedicine 2017, 12, 1985.

(16) Lindahl,; Abraham,; Hess,; van der Spoel, GROMACS 2019.6 Manual. 2020; https://doi.org/10.5281/zenodo.3685925.

(17) Promoting transparency and reproducibility in enhanced molecular simulations. Nature Methods 2019, 16, 670–673.

(18) Tribello, G. A.; Bonomi, M.; Branduardi, D.; Camilloni, C.; Bussi, G. PLUMED 2: New feathers for an old bird. Computer Physics Communications 2014, 185, 604–613.

(19) Bonomi, M.; Branduardi, D.; Bussi, G.; Camilloni, C.; Provasi, D.; Raiteri, P.; Donadio, D.; Marinelli, F.; Pietrucci, F.; Broglia, R. A., et al. PLUMED: A portable plugin for free-energy calculations with molecular dynamics. Computer Physics Communications 2009, 180, 1961–1972.

(20) Horta, B. A.; Merz, P. T.; Fuchs, P. F.; Dolenc, J.; Riniker, S.; Hunenberger, P. H. A GROMOS-compatible force field for small organic molecules in the condensed phase: The 2016H66 parameter set. Journal of Chemical Theory and Computation 2016, 12, 3825–3850.

(21) Brooks, B. R.; Brooks III, C. L.; Mackerell Jr, A. D.; Nilsson, L.; Petrella, R. J.; Roux, B.; Won, Y.; Archontis, G.; Bartels, C.; Boresch, S., et al. CHARMM: the biomolecular simulation program. Journal of Computational Chemistry 2009, 30, 1545–1614.

(22) Lear, S.; Cobb, S. L. Pep-Calc. com: a set of web utilities for the calculation of peptide and peptoid properties and automatic mass spectral peak assignment. Journal of Computer-Aided Molecular Design 2016, 30, 271–277.

(23) Rostkowski, M.; Olsson, M. H.; Søndergaard, C. R.; Jensen, J. H. Graphical analysis of pH-dependent properties of proteins predicted using PROPKA. BMC Structural Biology 2011, 11, 1–6.

(24) Samantray, S.; Schumann, W.; Illig, A.-M.; Carballo-Pacheco, M.; Paul, A.; Barz, B.; Strodel, B. Computer Simulations of Aggregation of Proteins and Peptides; Springer, 2022; pp 235–279.

(25) Hess, B.; Bekker, H.; Berendsen, H. J.; Fraaije, J. G. LINCS: a linear constraint solver for molecular simulations. Journal of Computational Chemistry 1997, 18, 1463–1472.

(26) Darden, T.; York, D.; Pedersen, L. Particle mesh Ewald: An Nlog(N) method for Ewald sums in large systems. The Journal of Chemical Physics 1993, 98, 10089–10092.

(27) Bussi, G.; Donadio, D.; Parrinello, M. Canonical sampling through velocity rescaling. The Journal of Chemical Physics 2007, 126, 014101.

(28) Parrinello, M.; Rahman, A. Polymorphic transitions in single crystals: A new molecular dynamics method. Journal of Applied Physics 1981, 52, 7182–7190.

(29) Singh, G.; Brovchenko, I.; Oleinikova, A.; Winter, R. Peptide aggregation in finite systems. Biophysical Journal 2008, 95, 3208–3221.

(30) Hamsici, S.; White, A. D.; Acar, H. Peptide framework for screening the effects of amino acids on assembly. Science Advances 2022, 8, eabj0305.

(31) Yang, Z. Hofmeister effects: an explanation for the impact of ionic liquids on biocatalysis. Journal of Biotechnology 2009, 144, 12–22.

(32) Zhang, F.; Skoda, M.; Jacobs, R.; Zorn, S.; Martin, R. A.; Martin, C.; Clark, G.; Weggler, S.; Hildebrandt, A.; Kohlbacher, O., et al. Reentrant condensation of proteins in solution induced by multivalent counterions. Physical Review Letters 2008, 101, 148101.

(33) Kuroda, Y.; Suenaga, A.; Sato, Y.; Kosuda, S.; Taiji, M. All-atom molecular dynamics analysis of multi-peptide systems reproduces peptide solubility in line with experimental observations. Scientific Reports 2016, 6, 19479.

(34) Bragina, M. E.; Cirauqui, N.; Dal Peraro, M.; Santos, R. A.; Fraga-Silva, R. A.; Stergiopulos, N. Abstract P250: Unveiling Binding Pocket Structure Of Mas Receptor And Its Interaction With Angiotensin-(1-7). Hypertension 2018, 72, AP250–AP250.

(35) Durik, M.; van Veghel, R.; Kuipers, A.; Rink, R.; Haas Jimoh Akanbi, M.; Moll, G.; Danser, A.; Roks, A. J. The effect of the thioether-bridged, stabilized Angiotensin-(1–7) analogue cyclic ang-(1–7) on cardiac remodeling and endothelial function in rats with myocardial infarction. International Journal of Hypertension 2012, 2012.

