## Supplementary material for "Atomistic molecular insight on Angiotensin-(1-7) interpeptide interactions": Source code: bibliography.bib

%%%%%% new
@book{avanti2012innovative,
  title={Innovative strategies for stabilization of therapeutic peptides in aqueous formulations},
  author={Avanti, Christina},
  year={2012}
}

@article{wang2022therapeutic,
  title={Therapeutic peptides: current applications and future directions},
  author={Wang, Lei and Wang, Nanxi and Zhang, Wenping and Cheng, Xurui and Yan, Zhibin and Shao, Gang and Wang, Xi and Wang, Rui and Fu, Caiyun},
  journal={Signal Transduction and Targeted Therapy},
  volume={7},
  number={1},
  pages={1--27},
  year={2022},
  publisher={Nature Publishing Group}
}

@article{lau2018therapeutic,
  title={Therapeutic peptides: Historical perspectives, current development trends, and future directions},
  author={Lau, Jolene L and Dunn, Michael K},
  journal={Bioorganic \& Medicinal Chemistry},
  volume={26},
  number={10},
  pages={2700--2707},
  year={2018},
  publisher={Elsevier}
}

@article{zapadka2017factors,
  title={Factors affecting the physical stability (aggregation) of peptide therapeutics},
  author={Zapadka, Karolina L and Becher, Frederik J and Gomes dos Santos, AL and Jackson, Sophie E},
  journal={Interface Focus},
  volume={7},
  number={6},
  pages={20170030},
  year={2017},
  publisher={The Royal Society}
}

@article{iusuf2008angiotensin,
  title={Angiotensin-(1--7): pharmacological properties and pharmacotherapeutic perspectives},
  author={Iusuf, Dilek and Henning, Robert H and van Gilst, Wiek H and Roks, Anton JM},
  journal={European Journal of Pharmacology},
  volume={585},
  number={2-3},
  pages={303--312},
  year={2008},
  publisher={Elsevier}
}

@article{khajehpour2021targeting,
  title={Targeting the protective arm of the renin-angiotensin system: Focused on angiotensin-(1--7)},
  author={Khajehpour, Sana and Aghazadeh-Habashi, Ali},
  journal={Journal of Pharmacology and Experimental Therapeutics},
  volume={377},
  number={1},
  pages={64--74},
  year={2021},
  publisher={ASPET}
}

@article{lula2007study,
  title={Study of angiotensin-(1--7) vasoactive peptide and its $\beta$-cyclodextrin inclusion complexes: complete sequence-specific NMR assignments and structural studies},
  author={Lula, Ivana and Denadai, {\^A}ngelo L and Resende, Jarbas M and de Sousa, Frederico B and de Lima, Guilherme F and Pilo-Veloso, Dorila and Heine, Thomas and Duarte, H{\'e}lio A and Santos, Robson AS and Sinisterra, Rub{\'e}n D},
  journal={Peptides},
  volume={28},
  number={11},
  pages={2199--2210},
  year={2007},
  publisher={Elsevier}
}

@article{chi2022unveiling,
  title={Unveiling the G4-PAMAM capacity to bind and protect Ang-(1-7) bioactive peptide by molecular dynamics simulations},
  author={Chi, L and Asgharpour, Somayeh and Correa-Basurto, Jos{\'e} and Bandala, Cindy Rodr{\'\i}guez and Mart{\'\i}nez-Archundia, Marlet},
  journal={Journal of Computer-aided Molecular Design},
  pages={1--23},
  year={2022},
  publisher={Springer}
}

@article{russ2019stabilization,
  title={Stabilization of Angiotensin-(1-7) in Cardioprotective Solutions},
  author={Russ, Miriam and Jauk, Susanne and Wintersteiger, Reinhold and Gesslbauer, Bernd and Greilberger, Joachim and Andr{\"a}, Michaela and Ortner, Astrid},
  journal={International Journal of Peptide Research and Therapeutics},
  volume={25},
  number={4},
  pages={1271--1278},
  year={2019},
  publisher={Springer}
}

@misc{sigmadatasheet,
  title={Angiotensin Fragment 1-7 acetate salt hydrate},
  author={Merck KGaA},
  year={2022},
  url={https://www.sigmaaldrich.com/MX/es/product/sigma/a9202}
}

@misc{apexbio,
  title={Angiotensin (1-7)
},
  author={ApexBio Technology},
  year={2013},
  url={https://www.apexbt.com/angiotensin-1-7.html#cite}
}

%%%%%%%%%%%%%%%%%%%%%%%%%%%%%%%%%%%%%%%%%%%%%%%%

@article{van2005gromacs,
  title={GROMACS: fast, flexible, and free},
  author={Van Der Spoel, David and Lindahl, Erik and Hess, Berk and Groenhof, Gerrit and Mark, Alan E and Berendsen, Herman JC},
  journal={Journal of Computational Chemistry},
  volume={26},
  number={16},
  pages={1701--1718},
  year={2005},
  publisher={Wiley Online Library}
}

@article{horta2016gromos,
  title={A GROMOS-compatible force field for small organic molecules in the condensed phase: The 2016H66 parameter set},
  author={Horta, Bruno AC and Merz, Pascal T and Fuchs, Patrick FJ and Dolenc, Jozica and Riniker, Sereina and Hunenberger, Philippe H},
  journal={Journal of Chemical Theory and Computation},
  volume={12},
  number={8},
  pages={3825--3850},
  year={2016},
  publisher={ACS Publications}
}

@article{lear2016pep,
  title={Pep-Calc. com: a set of web utilities for the calculation of peptide and peptoid properties and automatic mass spectral peak assignment},
  author={Lear, Sam and Cobb, Steven L},
  journal={Journal of Computer-Aided Molecular Design},
  volume={30},
  number={3},
  pages={271--277},
  year={2016},
  publisher={Springer}
}

@article{ramos2021pypolybuilder,
  title={pyPolyBuilder: Automated Preparation of Molecular Topologies and Initial Configurations for Molecular Dynamics Simulations of Arbitrary Supramolecules},
  author={Ramos, Mayk C and Quoika, Patrick K and Horta, Vitor AC and Dias, Douglas M and Costa, Elan G and do Amaral, Jorge LM and Ribeiro, Luigi M and Liedl, Klaus R and Horta, Bruno AC},
  journal={Journal of Chemical Information and Modeling},
  volume={61},
  number={4},
  pages={1539--1544},
  year={2021},
  publisher={ACS Publications},
}

@article{cakara2003microscopic,
  title={Microscopic protonation equilibria of poly (amidoamine) dendrimers from macroscopic titrations},
  author={Cakara, Dusko and Kleimann, J{\"o}rg and Borkovec, Michal},
  journal={Macromolecules},
  volume={36},
  number={11},
  pages={4201--4207},
  year={2003},
  publisher={ACS Publications}
}

@article{hess1997lincs,
  title={LINCS: a linear constraint solver for molecular simulations},
  author={Hess, Berk and Bekker, Henk and Berendsen, Herman JC and Fraaije, Johannes GEM},
  journal={Journal of Computational Chemistry},
  volume={18},
  number={12},
  pages={1463--1472},
  year={1997},
  publisher={Wiley Online Library}
}

@article{darden1993particle,
  title={Particle mesh Ewald: An Nlog(N) method for Ewald sums in large systems},
  author={Darden, Tom and York, Darrin and Pedersen, Lee},
  journal={The Journal of Chemical Physics},
  volume={98},
  number={12},
  pages={10089--10092},
  year={1993},
  publisher={American Institute of Physics}
}

@article{ramos2019molecular,
  title={Molecular dynamics simulations of PAMAM and PPI dendrimers using the GROMOS-Compatible 2016H66 Force Field},
  author={Ramos, Mayk C and Horta, Vitor AC and Horta, Bruno AC},
  journal={Journal of Chemical Information and Modeling},
  volume={59},
  number={4},
  pages={1444--1457},
  year={2019},
  publisher={ACS Publications}
}

@article{maingi2012dendrimer,
  title={Dendrimer building toolkit: Model building and characterization of various dendrimer architectures},
  author={Maingi, Vishal and Jain, Vaibhav and Bharatam, Prasad V and Maiti, Prabal K},
  journal={Journal of Computational Chemistry},
  volume={33},
  number={25},
  pages={1997--2011},
  year={2012},
  publisher={Wiley Online Library}
}

@article{caballero2013association,
  title={Association of nicotinic acid with a poly (amidoamine) dendrimer studied by molecular dynamics simulations},
  author={Caballero, Julio and Poblete, Horacio and Navarro, Cristell and Alzate-Morales, Jans H},
  journal={Journal of Molecular Graphics and Modelling},
  volume={39},
  pages={71--78},
  year={2013},
  publisher={Elsevier}
}

@article{porcar2008structural,
  title={Structural investigation of PAMAM dendrimers in aqueous solutions using small-angle neutron scattering: effect of generation},
  author={Porcar, Lionel and Liu, Yun and Verduzco, Rafael and Hong, Kunlun and Butler, Paul D and Magid, Linda J and Smith, Gregory S and Chen, Wei-Ren},
  journal={The Journal of Physical Chemistry B},
  volume={112},
  number={47},
  pages={14772--14778},
  year={2008},
  publisher={ACS Publications}
}

@article{rathgeber2002dynamics,
  title={Dynamics of star-burst dendrimers in solution in relation to their structural properties},
  author={Rathgeber, S and Monkenbusch, M and Kreitschmann, M and Urban, V and Brulet, A},
  journal={The Journal of Chemical Physics},
  volume={117},
  number={8},
  pages={4047--4062},
  year={2002},
  publisher={American Institute of Physics}
}

@article{badalkhani2017atomistic,
  title={Atomistic computer simulations on multi-loaded PAMAM dendrimers: A comparison of amine-and hydroxyl-terminated dendrimers},
  author={Badalkhani-Khamseh, Farideh and Ebrahim-Habibi, Azadeh and Hadipour, Nasser L},
  journal={Journal of Computer-Aided Molecular Design},
  volume={31},
  number={12},
  pages={1097--1111},
  year={2017},
  publisher={Springer}
}

@article{rostkowski2011graphical,
  title={Graphical analysis of pH-dependent properties of proteins predicted using PROPKA},
  author={Rostkowski, Micha{\l} and Olsson, Mats HM and S{\o}ndergaard, Chresten R and Jensen, Jan H},
  journal={BMC Structural Biology},
  volume={11},
  number={1},
  pages={1--6},
  year={2011},
  publisher={Springer}
}

@article{garcia2020host,
  title={Host-guest complexation of curcumin and coumarin 6 with PAMAM-OH: Insight from fluorescence spectroscopy and molecular dynamics simulations},
  author={Garc{\'\i}a, Yadiris and Jim{\'e}nez, Ver{\'o}nica A and Alderete, Joel B},
  journal={Journal of Luminescence},
  volume={222},
  pages={117182},
  year={2020},
  publisher={Elsevier}
}

@article{tanis2009association,
  title={Association of a weakly acidic anti-inflammatory drug (ibuprofen) with a poly (amidoamine) dendrimer as studied by molecular dynamics simulations},
  author={Tanis, I and Karatasos, K},
  journal={The Journal of Physical Chemistry B},
  volume={113},
  number={31},
  pages={10984--10993},
  year={2009},
  publisher={ACS Publications}
}

@article{daura1999peptide,
  title={Peptide folding: when simulation meets experiment},
  author={Daura, Xavier and Gademann, Karl and Jaun, Bernhard and Seebach, Dieter and Van Gunsteren, Wilfred F and Mark, Alan E},
  journal={Angewandte Chemie International Edition},
  volume={38},
  number={1-2},
  pages={236--240},
  year={1999},
  publisher={Wiley Online Library}
}

@article{trott2010autodock,
  title={AutoDock Vina: improving the speed and accuracy of docking with a new scoring function, efficient optimization, and multithreading},
  author={Trott, Oleg and Olson, Arthur J},
  journal={Journal of Computational Chemistry},
  volume={31},
  number={2},
  pages={455--461},
  year={2010},
  publisher={Wiley Online Library}
}

@article{capelli2019exhaustive,
  title={Exhaustive search of ligand binding pathways via volume-based metadynamics},
  author={Capelli, Riccardo and Carloni, Paolo and Parrinello, Michele},
  journal={The Journal of Physical Chemistry Letters},
  volume={10},
  number={12},
  pages={3495--3499},
  year={2019},
  publisher={ACS Publications}
}

@article{shylesh2022renin,
  title={Renin-angiotensin system modulators in COVID-19 patients with hypertension: friend or foe?},
  author={Shylesh CM, Shakhi and VS, Arya and SK, Kanthlal},
  journal={Clinical and Experimental Hypertension},
  volume={44},
  number={1},
  pages={1--10},
  year={2022},
  publisher={Taylor \& Francis}
}

@article{muslim2020treatment,
  title={Treatment options available for COVID-19 and an analysis on possible role of combination of rhACE2, angiotensin (1-7) and angiotensin (1-9) as effective therapeutic measure},
  author={Muslim, Shahnawaz and Nasrin, Nasrin and Alotaibi, Faisal Obaid and Prasad, Geeta and Singh, Shambhu Kumar and Alam, Izhar and Mustafa, Gulam},
  journal={SN Comprehensive Clinical Medicine},
  volume={2},
  number={10},
  pages={1761--1766},
  year={2020},
  publisher={Springer}
}

@article{soto2020countermeasure,
  title={Countermeasure and therapeutic: A (1--7) to treat acute respiratory distress syndrome due to COVID-19 infection},
  author={Soto, Maira and DiZerega, Gere and Rodgers, Kathleen E},
  journal={Journal of the Renin-Angiotensin-Aldosterone System},
  volume={21},
  number={4},
  pages={1470320320972018},
  year={2020},
  publisher={SAGE Publications Sage UK: London, England}
}

@article{imanpour2020angiotensin,
  title={Angiotensin 1-7: A novel strategy in COVID-19 Treatment},
  author={Imanpour, Hamed and Rezaee, Haleh and Nouri-Vaskeh, Masoud},
  journal={Advanced Pharmaceutical Bulletin},
  volume={10},
  number={4},
  pages={488},
  year={2020},
  publisher={Tabriz University of Medical Sciences}
}

@article{peiro2020substituting,
  title={Substituting angiotensin-(1-7) to prevent lung damage in SARS-CoV-2 infection?},
  author={Peir{\'o}, Concepci{\'o}n and Moncada, Salvador},
  journal={Circulation},
  volume={141},
  number={21},
  pages={1665--1666},
  year={2020},
  publisher={Am Heart Assoc}
}

@article{issa2021combination,
  title={Combination of angiotensin (1-7) agonists and convalescent plasma as a new strategy to overcome angiotensin converting enzyme 2 (ACE2) inhibition for the treatment of COVID-19},
  author={Issa, Hawraa and Eid, Ali H and Berry, Bassam and Takhviji, Vahideh and Khosravi, Abbas and Mantash, Sarah and Nehme, Rawan and Hallal, Rawan and Karaki, Hussein and Dhayni, Kawthar and others},
  journal={Frontiers in Medicine},
  volume={8},
  pages={278},
  year={2021},
  publisher={Frontiers}
}

@article{petty2009phase,
  title={Phase I and pharmacokinetic study of angiotensin-(1-7), an endogenous antiangiogenic hormone},
  author={Petty, W Jeffrey and Miller, Antonius A and McCoy, Thomas P and Gallagher, Patricia E and Tallant, E Ann and Torti, Frank M},
  journal={Clinical Cancer Research},
  volume={15},
  number={23},
  pages={7398--7404},
  year={2009},
  publisher={AACR}
}

@article{wester2017stabilization,
  title={Stabilization of Angiotensin-(1--7) by key substitution with a cyclic non-natural amino acid},
  author={Wester, Anita and Devocelle, Marc and Tallant, E and Chappell, Mark C and Gallagher, Patricia E and Paradisi, Francesca},
  journal={Amino Acids},
  volume={49},
  number={10},
  pages={1733--1742},
  year={2017},
  publisher={Springer}
}

@article{durik2012effect,
  title={The effect of the thioether-bridged, stabilized Angiotensin-(1--7) analogue cyclic ang-(1--7) on cardiac remodeling and endothelial function in rats with myocardial infarction},
  author={Durik, Matej and van Veghel, Richard and Kuipers, Anneke and Rink, Rick and Haas Jimoh Akanbi, Marijke and Moll, Gert and Danser, AH and Roks, Anton JM},
  journal={International Journal of Hypertension},
  volume={2012},
  year={2012},
  publisher={Hindawi}
}
@article{de2007study,
  title={Study of angiotensin-(1--7) vasoactive peptide and its b-cyclodextrin inclusion complexes: Complete sequence-specific NMR assignments and structural studies},
  author={de Lima, Guilherme F and Pilo-Veloso, Dorila and Heine, Thomas and Duarte, H{\'e}lio A and Santos, Robson AS and Sinisterra, Rub{\'e}n D},
  journal={Peptides},
  volume={28},
  pages={2199--2210},
  year={2007}
}

@article{marquez2017complex,
  title={The complex of PAMAM-OH dendrimer with Angiotensin (1--7) prevented the disuse-induced skeletal muscle atrophy in mice},
  author={M{\'a}rquez-Miranda, Valeria and Abrigo, Johanna and Rivera, Juan Carlos and Araya-Duran, Ingrid and Aravena, Javier and Simon, Felipe and Pacheco, Nicolas and Gonz{\'a}lez-Nilo, Fernando Danilo and Cabello-Verrugio, Claudio},
  journal={International Journal of Nanomedicine},
  volume={12},
  pages={1985},
  year={2017},
  publisher={Dove Press}
}

@article{rodriguez2019silico,
  title={In silico search, chemical characterization and immunogenic evaluation of amino-terminated G4-PAMAM-HIV peptide complexes using three-dimensional models of the HIV-1 gp120 protein},
  author={Rodr{\'\i}guez-Fonseca, Rolando Alberto and Bello, Martiniano and de Los Mu{\~n}oz-Fern{\'a}ndez, Mar{\'\i}a {\'A}ngeles and Jim{\'e}nez, Jos{\'e} Luis and Rojas-Hern{\'a}ndez, Sa{\'u}l and Fragoso-V{\'a}zquez, MJ and Guti{\'e}rrez-S{\'a}nchez, Mara and Rodrigues, Jo{\~a}o and Cayetano-Castro, N and Borja-Urby, R and others},
  journal={Colloids and Surfaces B: Biointerfaces},
  volume={177},
  pages={77--93},
  year={2019},
  publisher={Elsevier}
}
@article{bello2020complexation,
  title={Complexation of peptide epitopes with G4-PAMAM dendrimer through ligand diffusion molecular dynamic simulations},
  author={Bello, Martiniano and Rodr{\'\i}guez-Fonseca, Rolando Alberto and Correa-Basurto, Jos{\'e}},
  journal={Journal of Molecular Graphics and Modelling},
  volume={96},
  pages={107514},
  year={2020},
  publisher={Elsevier}
}

@article{shcharbin2014study,
  title={How to study dendrimers and dendriplexes III. Biodistribution, pharmacokinetics and toxicity in vivo},
  author={Shcharbin, Dzmitry and Janaszewska, Anna and Klajnert-Maculewicz, Barbara and Ziemba, Barbara and Dzmitruk, Volha and Halets, Inessa and Loznikova, Svetlana and Shcharbina, Natallia and Milowska, Katarzyna and Ionov, Maksim and others},
  journal={Journal of Controlled Release},
  volume={181},
  pages={40--52},
  year={2014},
  publisher={Elsevier}
}

@misc{doubledock,
  title={Double-docking-approach},
  author={L. América Chi},
  year={2022},
  url={https://github.com/AmericaChi/Double-docking-approach}
}

@article{bussi2007canonical,
  title={Canonical sampling through velocity rescaling},
  author={Bussi, Giovanni and Donadio, Davide and Parrinello, Michele},
  journal={The Journal of Chemical Physics},
  volume={126},
  number={1},
  pages={014101},
  year={2007},
  publisher={American Institute of Physics}
}

@article{parrinello1981polymorphic,
  title={Polymorphic transitions in single crystals: A new molecular dynamics method},
  author={Parrinello, Michele and Rahman, Aneesur},
  journal={Journal of Applied Physics},
  volume={52},
  number={12},
  pages={7182--7190},
  year={1981},
  publisher={American Institute of Physics}
}

@article{fox2018pamam,
  title={PAMAM dendrimer-cell membrane interactions},
  author={Fox, Laura J and Richardson, Robert M and Briscoe, Wuge H},
  journal={Advances in Colloid and Interface Science},
  volume={257},
  pages={1--18},
  year={2018},
  publisher={Elsevier}
}

@article{barducci2013free,
  title={Free-energy landscape of protein oligomerization from atomistic simulations},
  author={Barducci, Alessandro and Bonomi, Massimiliano and Prakash, Meher K and Parrinello, Michele},
  journal={Proceedings of the National Academy of Sciences},
  volume={110},
  number={49},
  pages={E4708--E4713},
  year={2013},
  publisher={National Acad Sciences}
}

@article{barducci2011metadynamics,
  title={Metadynamics},
  author={Barducci, Alessandro and Bonomi, Massimiliano and Parrinello, Michele},
  journal={Wiley Interdisciplinary Reviews: Computational Molecular Science},
  volume={1},
  number={5},
  pages={826--843},
  year={2011},
  publisher={Wiley Online Library}
}

@article{laio2008metadynamics,
  title={Metadynamics: a method to simulate rare events and reconstruct the free energy in biophysics, chemistry and material science},
  author={Laio, Alessandro and Gervasio, Francesco L},
  journal={Reports on Progress in Physics},
  volume={71},
  number={12},
  pages={126601},
  year={2008},
  publisher={IOP Publishing}
}

@article{barducci2008well,
  title={Well-tempered metadynamics: a smoothly converging and tunable free-energy method},
  author={Barducci, Alessandro and Bussi, Giovanni and Parrinello, Michele},
  journal={Physical Review Letters},
  volume={100},
  number={2},
  pages={020603},
  year={2008},
  publisher={APS}
}

@article{tiwary2015time,
  title={A time-independent free energy estimator for metadynamics},
  author={Tiwary, Pratyush and Parrinello, Michele},
  journal={The Journal of Physical Chemistry B},
  volume={119},
  number={3},
  pages={736--742},
  year={2015},
  publisher={ACS Publications}
}

@article{silva2001long,
  title={Long-lasting cardiovascular effects of liposome-entrapped angiotensin-(1-7) at the rostral ventrolateral medulla},
  author={Silva-Barcellos, Neila M and Fr{\'e}zard, Fr{\'e}d{\'e}ric and Caligiorne, Sordaini and Santos, Robson AS},
  journal={Hypertension},
  volume={38},
  number={6},
  pages={1266--1271},
  year={2001},
  publisher={Am Heart Assoc}
}

@article{kluskens2009angiotensin,
  title={Angiotensin-(1-7) with thioether bridge: an angiotensin-converting enzyme-resistant, potent angiotensin-(1-7) analog},
  author={Kluskens, Leon D and Nelemans, S Adriaan and Rink, Rick and de Vries, Louwe and Meter-Arkema, Anita and Wang, Yong and Walther, Thomas and Kuipers, Anneke and Moll, Gert N and Haas, Marijke},
  journal={Journal of Pharmacology and Experimental Therapeutics},
  volume={328},
  number={3},
  pages={849--854},
  year={2009},
  publisher={ASPET}
}

@misc{tijsma2007angiotensin,
  title={Angiotensin-(1-7) eluting polymer-coated medical device to reduce restenosis and improve endothelial cell function},
  author={Tijsma, Edze Jan and Driessen-Levels, Anita and Hendriks, Marc},
  year={2007},
  month=feb # "~13",
  publisher={Google Patents},
  note={US Patent 7,176,261}
}

@article{chappell1998metabolism,
  title={Metabolism of angiotensin-(1--7) by angiotensin-converting enzyme},
  author={Chappell, Mark C and Pirro, Nancy T and Sykes, Angela and Ferrario, Carlos M},
  journal={Hypertension},
  volume={31},
  number={1},
  pages={362--367},
  year={1998},
  publisher={Am Heart Assoc}
}


@article{yamada1998converting,
  title={Converting enzyme determines plasma clearance of angiotensin-(1--7)},
  author={Yamada, Kazuo and Iyer, Shridhar N and Chappell, Mark C and Ganten, Detlev and Ferrario, Carlos M},
  journal={Hypertension},
  volume={32},
  number={3},
  pages={496--502},
  year={1998},
  publisher={Am Heart Assoc}
}

@article{pavan2013combining,
  title={Combining metadynamics simulation and experiments to characterize dendrimers in solution},
  author={Pavan, Giovanni M and Barducci, Alessandro and Albertazzi, Lorenzo and Parrinello, Michele},
  journal={Soft Matter},
  volume={9},
  number={9},
  pages={2593--2597},
  year={2013},
  publisher={Royal Society of Chemistry}
}

@article{clayton2022peptide,
  title={Peptide Dynamics and Metadynamics: Leveraging Enhanced Sampling Molecular Dynamics to Robustly Model Long-Timescale Transitions.},
  author={Clayton, Joseph and Baweja, Lokesh and Wereszczynski, Jeff},
  journal={Methods in Molecular Biology (Clifton, NJ)},
  volume={2405},
  pages={151--167},
  year={2022}
}

@article{ballauff2004dendrimers,
  title={Dendrimers in solution: insight from theory and simulation},
  author={Ballauff, Matthias and Likos, Christos N},
  journal={Angewandte Chemie International Edition},
  volume={43},
  number={23},
  pages={2998--3020},
  year={2004},
  publisher={Wiley Online Library}
}

@misc{covidtrial,
  title={Use of Angiotensin-(1-7) in COVID-19},
  author={ClinicalTrials.gov},
  year={2022},
  url={https://clinicaltrials.gov/ct2/show/NCT04633772}
}

@article{bader2018meaning,
  title={The meaning of Mas},
  author={Bader, Michael and Alenina, Natalia and Young, Dallan and Santos, Robson AS and Touyz, Rhian M},
  journal={Hypertension},
  volume={72},
  number={5},
  pages={1072--1075},
  year={2018},
  publisher={Am Heart Assoc}
}

@article{janaszewska2019cytotoxicity,
  title={Cytotoxicity of dendrimers},
  author={Janaszewska, Anna and Lazniewska, Joanna and Trzepi{\'n}ski, Przemys{\l}aw and Marcinkowska, Monika and Klajnert-Maculewicz, Barbara},
  journal={Biomolecules},
  volume={9},
  number={8},
  pages={330},
  year={2019},
  publisher={MDPI}
}

@article{otto2018poly,
  title={Poly (amidoamine) dendrimers as a pharmaceutical excipient. Are we there yet?},
  author={Otto, Daniel P and De Villiers, Melgardt M},
  journal={Journal of Pharmaceutical Sciences},
  volume={107},
  number={1},
  pages={75--83},
  year={2018},
  publisher={Elsevier}
}

@article{li2022safety,
  title={Safety Challenges and Application Strategies for the Use of Dendrimers in Medicine},
  author={Li, Xiang and Naeem, Abid and Xiao, Shanghua and Hu, Lei and Zhang, Jing and Zheng, Qin},
  journal={Pharmaceutics},
  volume={14},
  number={6},
  pages={1292},
  year={2022},
  publisher={Multidisciplinary Digital Publishing Institute}
}

@article{agashe2006investigations,
  title={Investigations on the toxicological profile of functionalized fifth-generation poly (propylene imine) dendrimer},
  author={Agashe, Hrushikesh B and Dutta, Tathagata and Garg, Minakshi and Jain, NK},
  journal={Journal of Pharmacy and Pharmacology},
  volume={58},
  number={11},
  pages={1491--1498},
  year={2006},
  publisher={Wiley Online Library}
}

@article{daliri2018current,
  title={Current trends and perspectives of bioactive peptides},
  author={Daliri, Eric Banan-Mwine and Lee, Byong H and Oh, Deog H},
  journal={Critical Reviews in Food Science and Nutrition},
  volume={58},
  number={13},
  pages={2273--2284},
  year={2018},
  publisher={Taylor \& Francis}
}

@article{fosgerau2015peptide,
  title={Peptide therapeutics: current status and future directions},
  author={Fosgerau, Keld and Hoffmann, Torsten},
  journal={Drug Discovery Today},
  volume={20},
  number={1},
  pages={122--128},
  year={2015},
  publisher={Elsevier}
}

@article{armiento2020peptide,
  title={Peptide-Based Molecular Strategies To Interfere with Protein Misfolding, Aggregation, and Cell Degeneration},
  author={Armiento, Valentina and Spanopoulou, Anna and Kapurniotu, Aphrodite},
  journal={Angewandte Chemie International Edition},
  volume={59},
  number={9},
  pages={3372--3384},
  year={2020},
  publisher={Wiley Online Library}
}

@misc{lindahl_2020_3685925,
  author       = {Lindahl and
                  Abraham and
                  Hess and
                  van der Spoel},
  title        = {GROMACS 2019.6 Manual},
  month        = feb,
  year         = 2020,
  publisher    = {Zenodo},
  version      = {2019.6},
  doi          = {10.5281/zenodo.3685925},
  url          = {https://doi.org/10.5281/zenodo.3685925}
}

@article{plumed2019promoting,
  title={Promoting transparency and reproducibility in enhanced molecular simulations},
  journal={Nature Methods},
  volume={16},
  number={8},
  pages={670--673},
  year={2019},
  publisher={Nature Publishing Group US New York}
}

@article{tribello2014plumed,
  title={PLUMED 2: New feathers for an old bird},
  author={Tribello, Gareth A and Bonomi, Massimiliano and Branduardi, Davide and Camilloni, Carlo and Bussi, Giovanni},
  journal={Computer Physics Communications},
  volume={185},
  number={2},
  pages={604--613},
  year={2014},
  publisher={Elsevier}
}

@article{bonomi2009plumed,
  title={PLUMED: A portable plugin for free-energy calculations with molecular dynamics},
  author={Bonomi, Massimiliano and Branduardi, Davide and Bussi, Giovanni and Camilloni, Carlo and Provasi, Davide and Raiteri, Paolo and Donadio, Davide and Marinelli, Fabrizio and Pietrucci, Fabio and Broglia, Ricardo A and others},
  journal={Computer Physics Communications},
  volume={180},
  number={10},
  pages={1961--1972},
  year={2009},
  publisher={Elsevier}
}

@article{brooks2009charmm,
  title={CHARMM: the biomolecular simulation program},
  author={Brooks, Bernard R and Brooks III, Charles L and Mackerell Jr, Alexander D and Nilsson, Lennart and Petrella, Robert J and Roux, Beno{\^\i}t and Won, Youngdo and Archontis, Georgios and Bartels, Christian and Boresch, Stefan and others},
  journal={Journal of Computational Chemistry},
  volume={30},
  number={10},
  pages={1545--1614},
  year={2009},
  publisher={Wiley Online Library}
}

@incollection{samantray2022molecular,
  title={Molecular dynamics simulations of protein aggregation: protocols for simulation setup and analysis with Markov state models and transition networks},
  author={Samantray, Suman and Schumann, Wibke and Illig, Alexander-Maurice and Carballo-Pacheco, Martin and Paul, Arghadwip and Barz, Bogdan and Strodel, Birgit},
  booktitle={Computer Simulations of Aggregation of Proteins and Peptides},
  pages={235--279},
  year={2022},
  publisher={Springer}
}

@article{singh2008peptide,
  title={Peptide aggregation in finite systems},
  author={Singh, Gurpreet and Brovchenko, Ivan and Oleinikova, Alla and Winter, Roland},
  journal={Biophysical Journal},
  volume={95},
  number={7},
  pages={3208--3221},
  year={2008},
  publisher={Elsevier}
}

@article{yang2009hofmeister,
  title={Hofmeister effects: an explanation for the impact of ionic liquids on biocatalysis},
  author={Yang, Zhen},
  journal={Journal of Biotechnology},
  volume={144},
  number={1},
  pages={12--22},
  year={2009},
  publisher={Elsevier}
}

@article{hamsici2022peptide,
  title={Peptide framework for screening the effects of amino acids on assembly},
  author={Hamsici, Seren and White, Andrew D and Acar, Handan},
  journal={Science Advances},
  volume={8},
  number={3},
  pages={eabj0305},
  year={2022},
  publisher={American Association for the Advancement of Science}
}

@article{kuroda2016all,
  title={All-atom molecular dynamics analysis of multi-peptide systems reproduces peptide solubility in line with experimental observations},
  author={Kuroda, Yutaka and Suenaga, Atsushi and Sato, Yuji and Kosuda, Satoshi and Taiji, Makoto},
  journal={Scientific Reports},
  volume={6},
  number={1},
  pages={19479},
  year={2016},
  publisher={Nature Publishing Group UK London}
}

@article{zhang2008reentrant,
  title={Reentrant condensation of proteins in solution induced by multivalent counterions},
  author={Zhang, Fan and Skoda, MWA and Jacobs, RMJ and Zorn, S and Martin, Richard A and Martin, CM and Clark, GF and Weggler, S and Hildebrandt, A and Kohlbacher, O and others},
  journal={Physical Review Letters},
  volume={101},
  number={14},
  pages={148101},
  year={2008},
  publisher={APS}
}

@article{bragina2018abstract,
  title={Abstract P250: Unveiling Binding Pocket Structure Of Mas Receptor And Its Interaction With Angiotensin-(1-7)},
  author={Bragina, Maiia E and Cirauqui, Nuria and Dal Peraro, Matteo and Santos, Robson A and Fraga-Silva, Rodrigo A and Stergiopulos, Nikos},
  journal={Hypertension},
  volume={72},
  number={Suppl\_1},
  pages={AP250--AP250},
  year={2018},
  publisher={Am Heart Assoc}
}
