## Supplementary figures and images for "Atomistic molecular insight on Angiotensin-(1-7) interpeptide interactions"

### ang-secondary.png

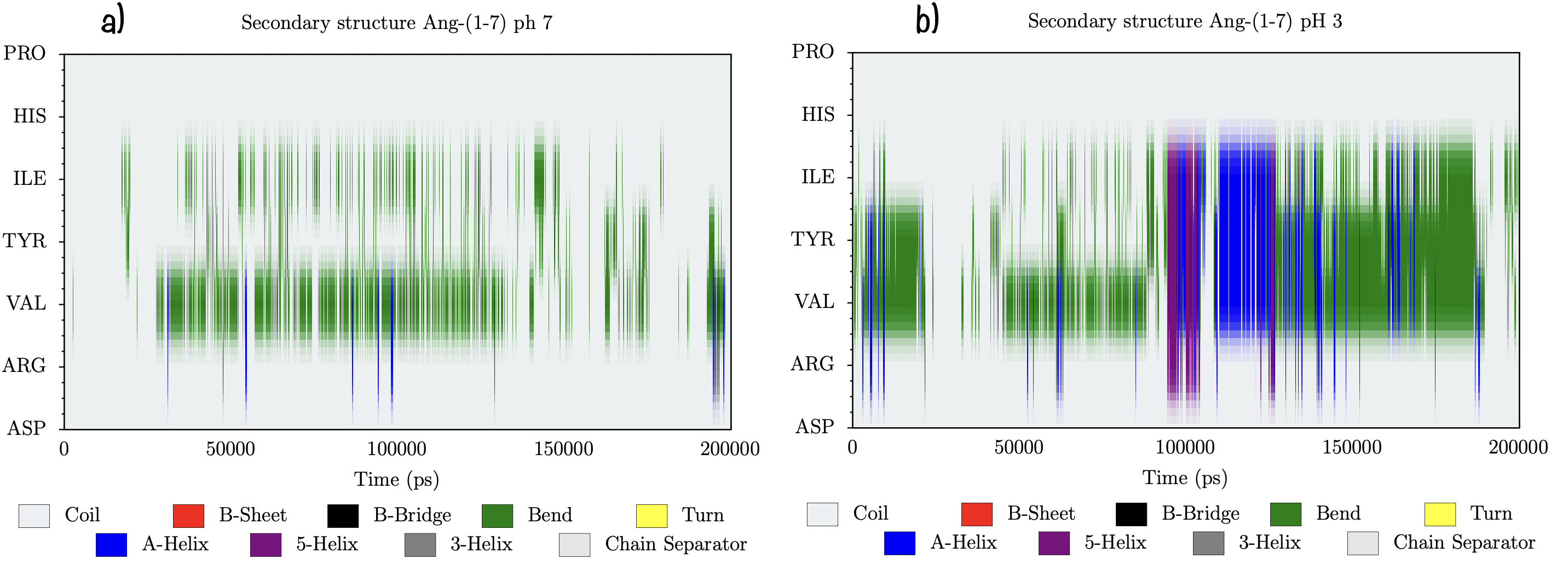

### cavity.png

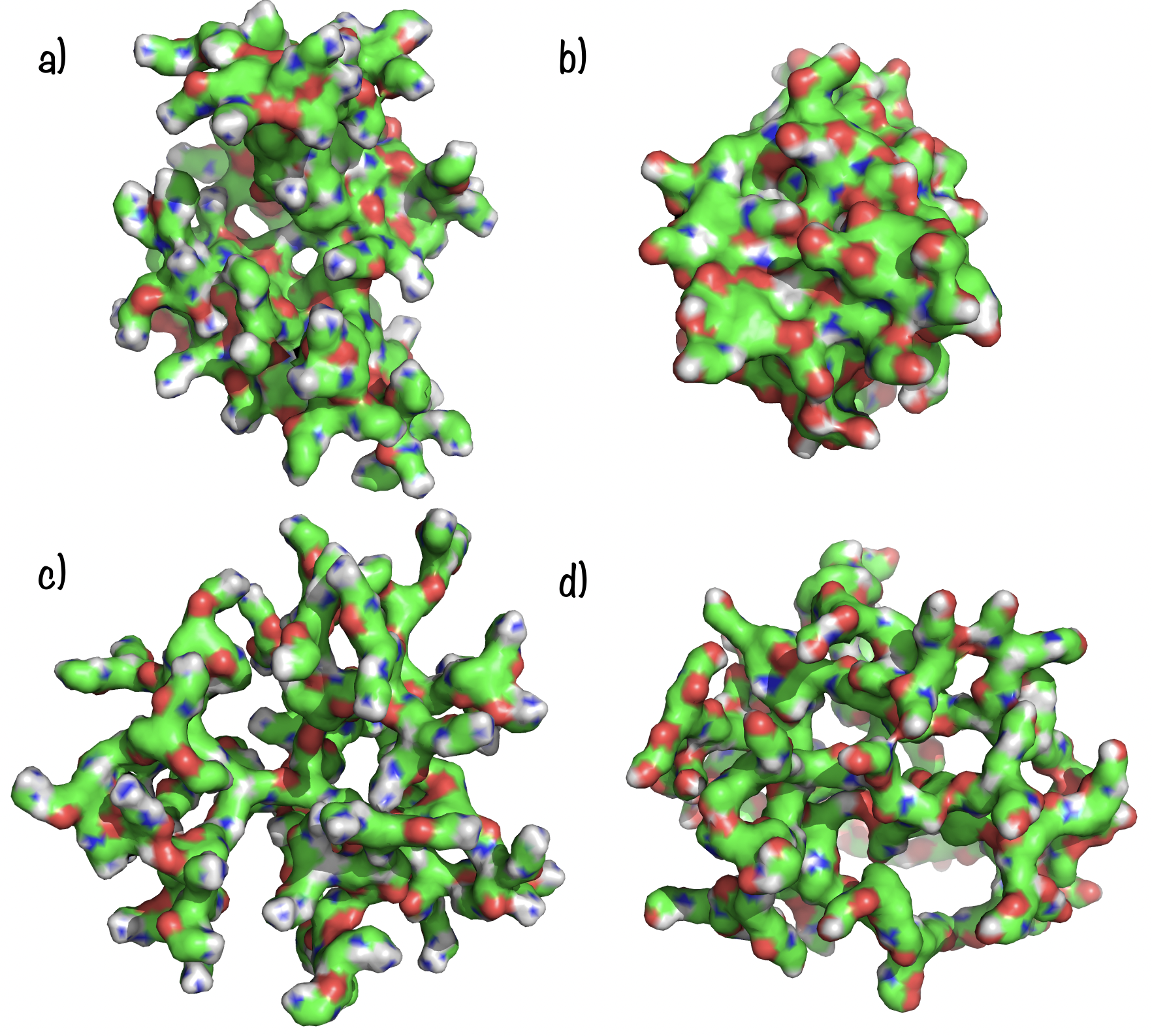

### clusterAnalysis.pdf

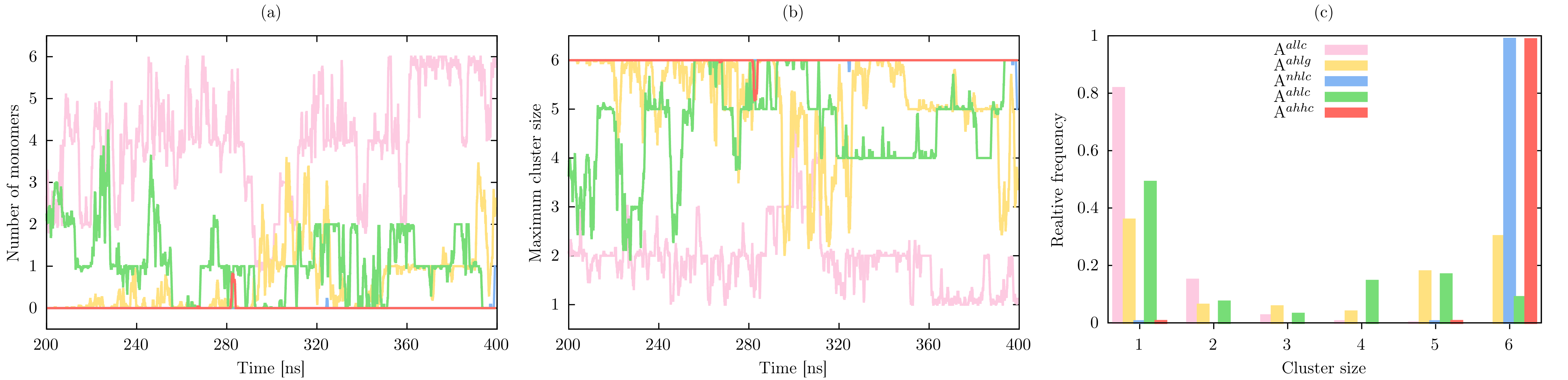

### clusters.pdf

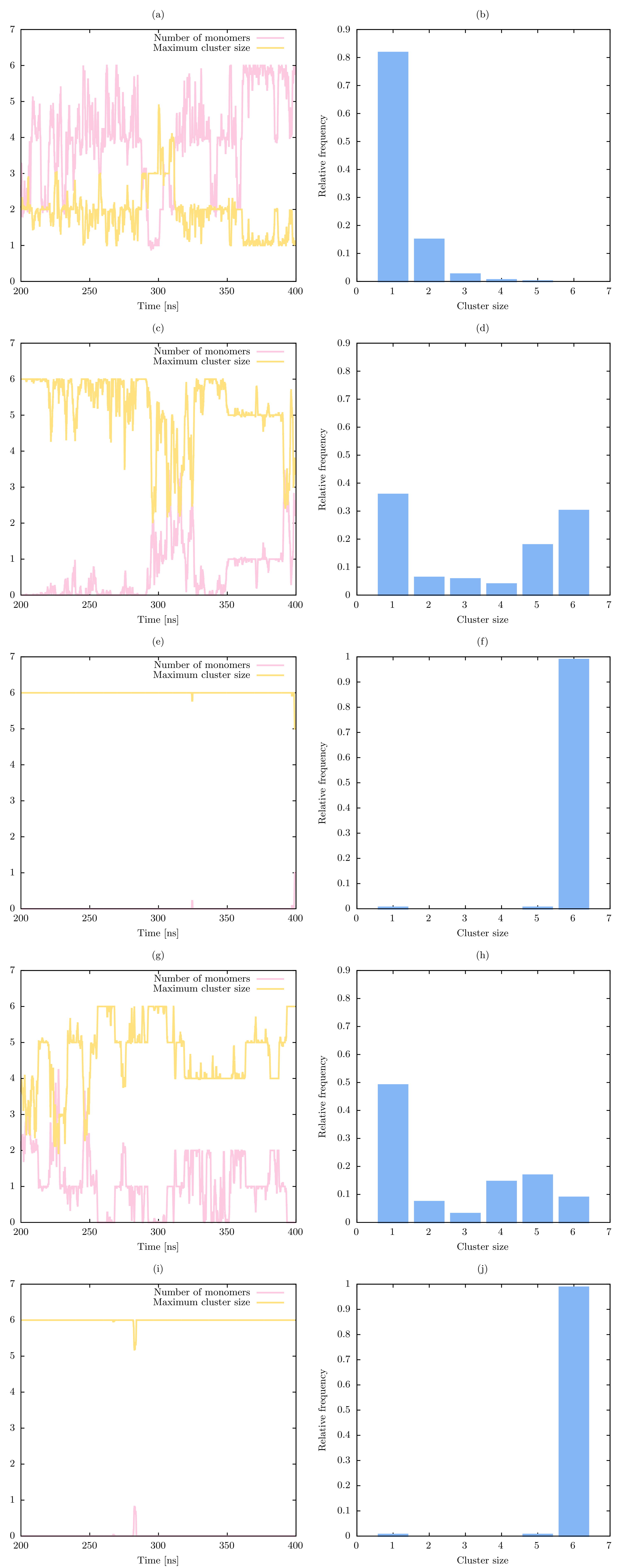

### comnh-properties-1.png

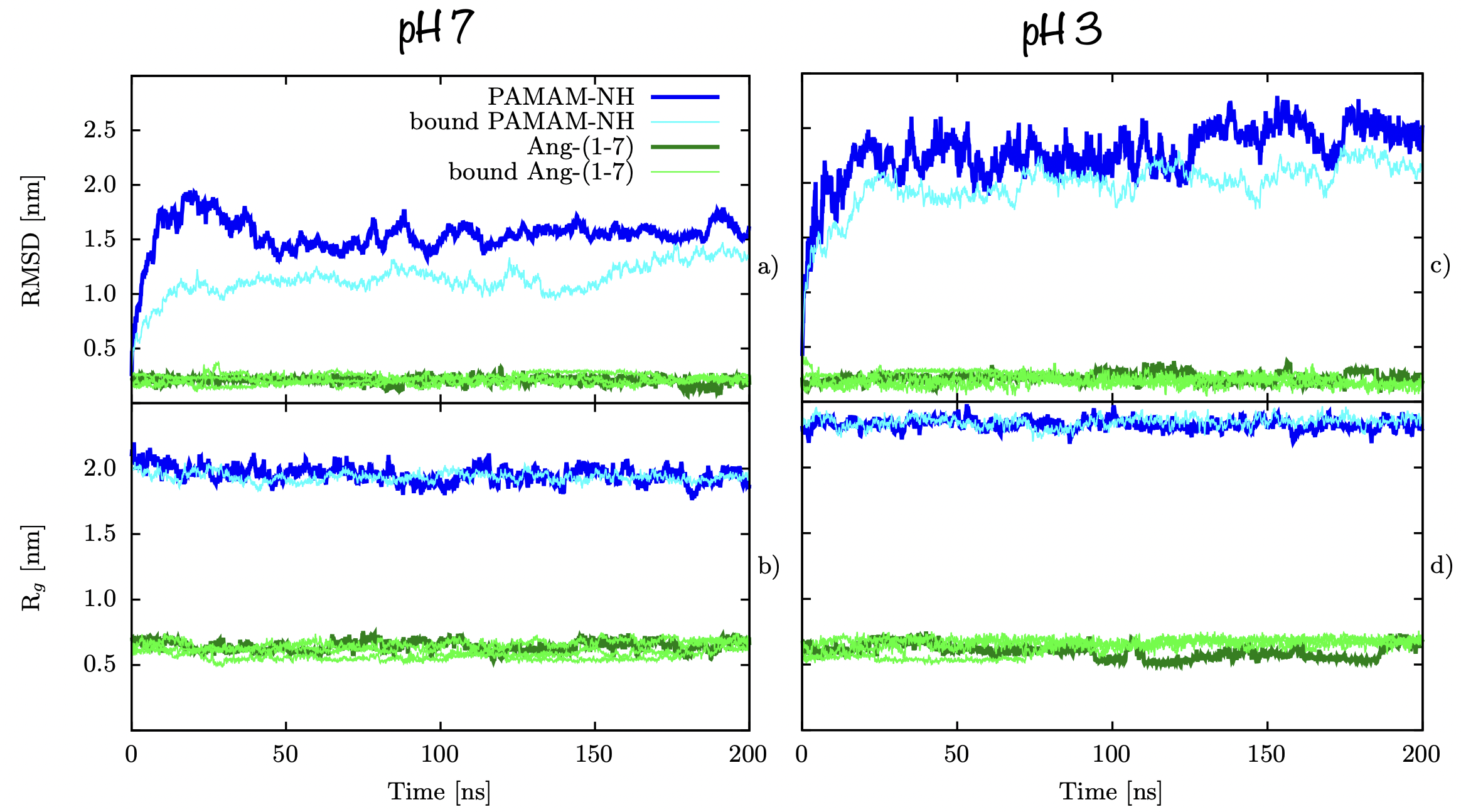

### comnh-properties-2.png

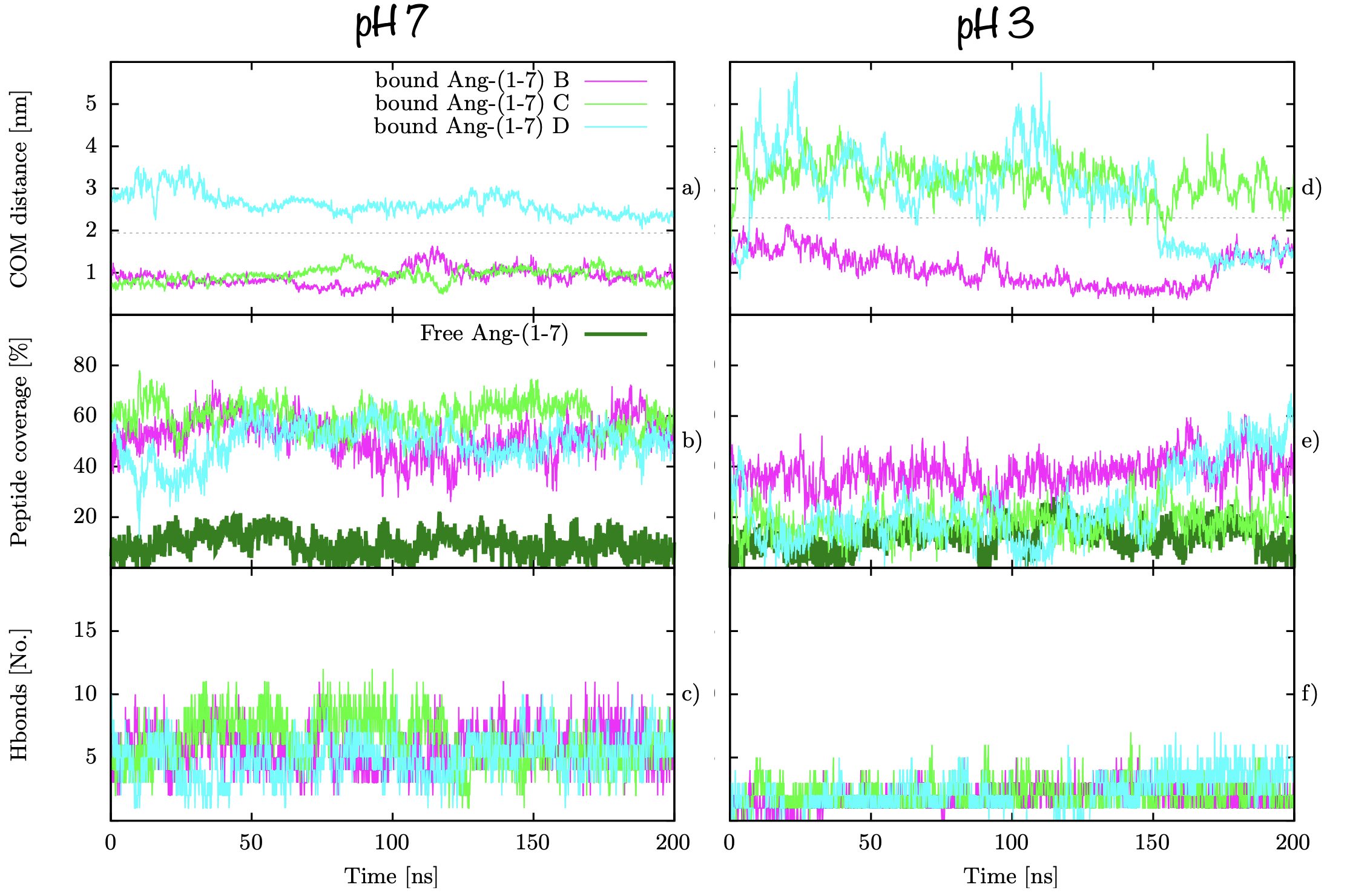

### comoh2-properties-2.png

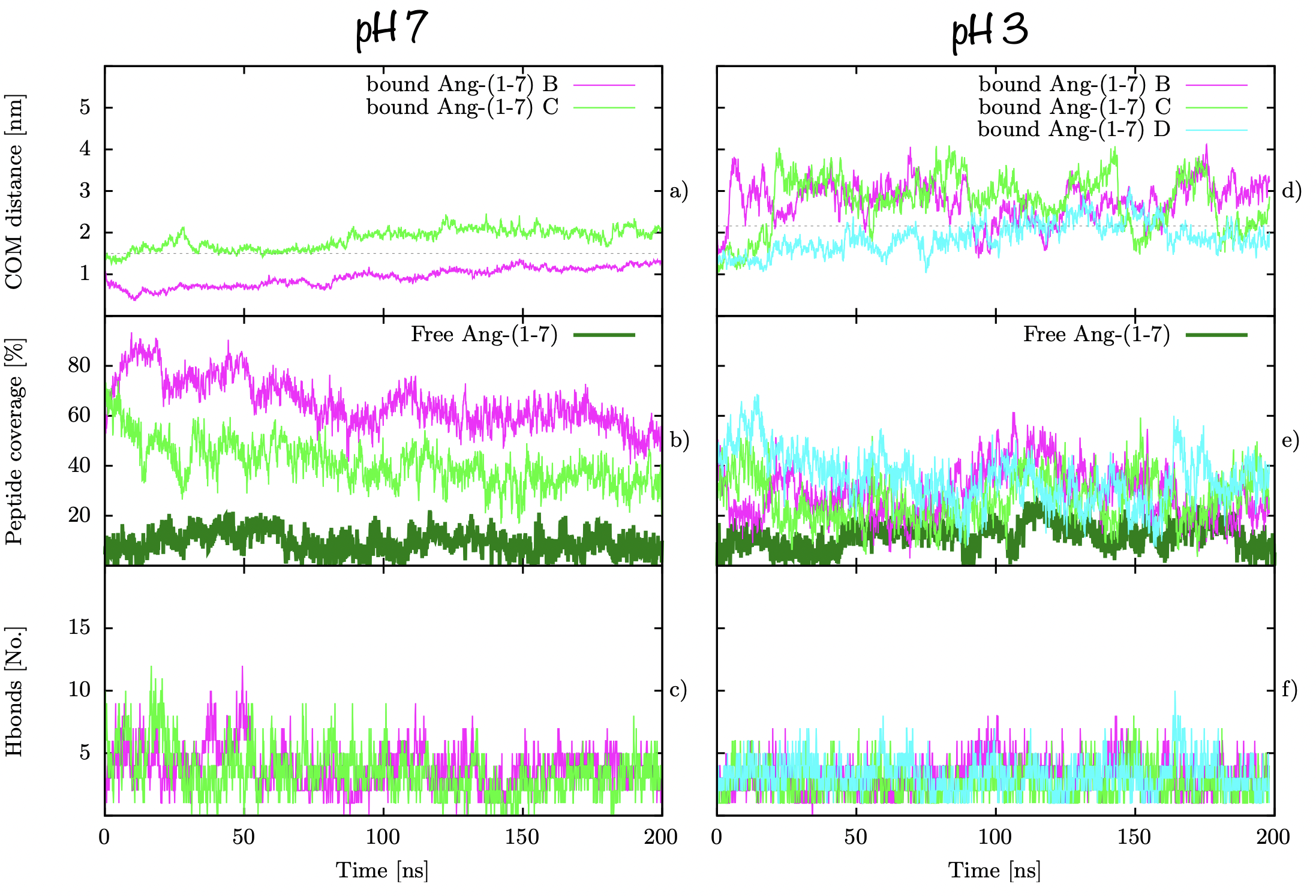

### comoh-properties-1.png

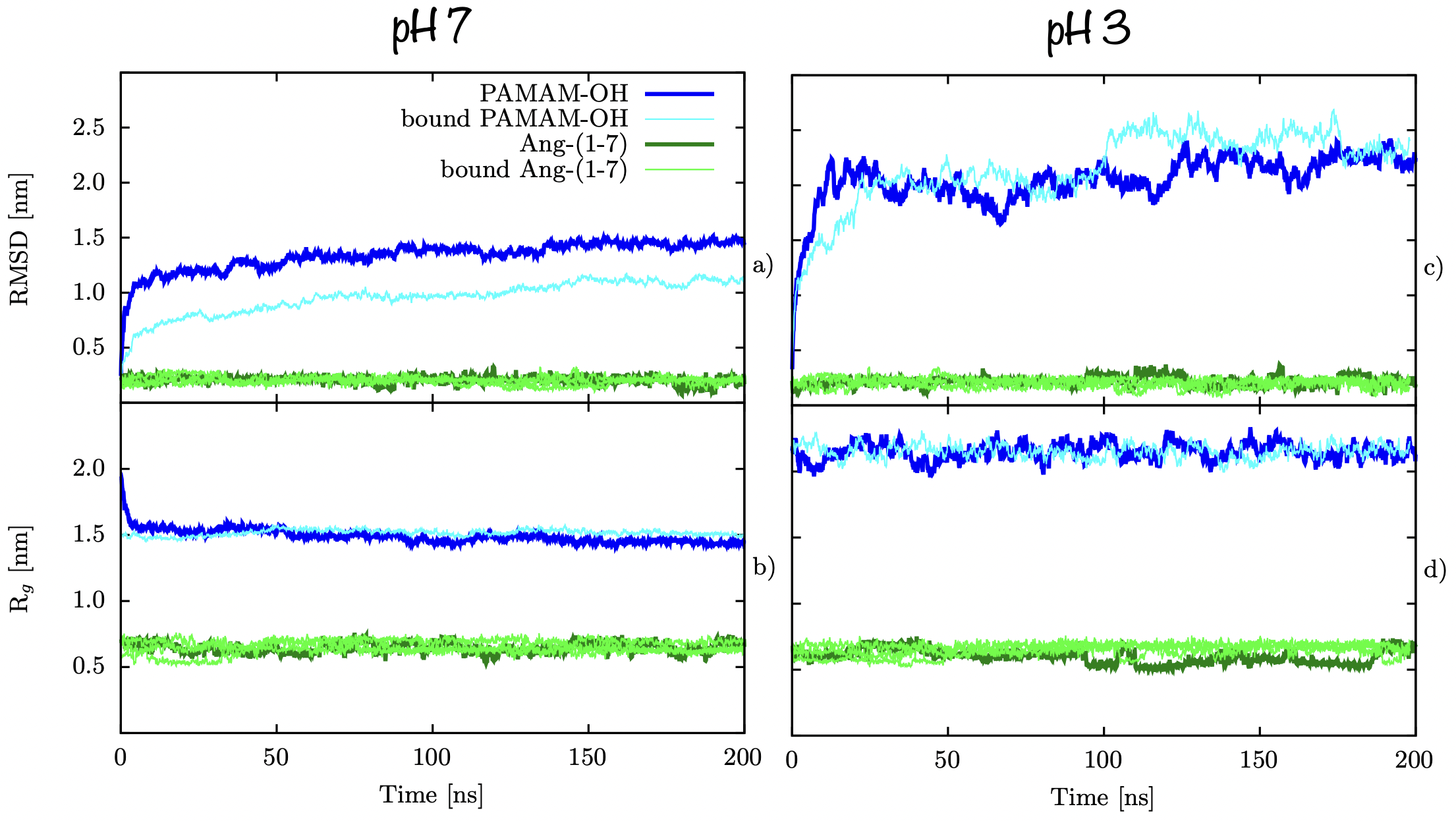

### comoh-properties-2.png

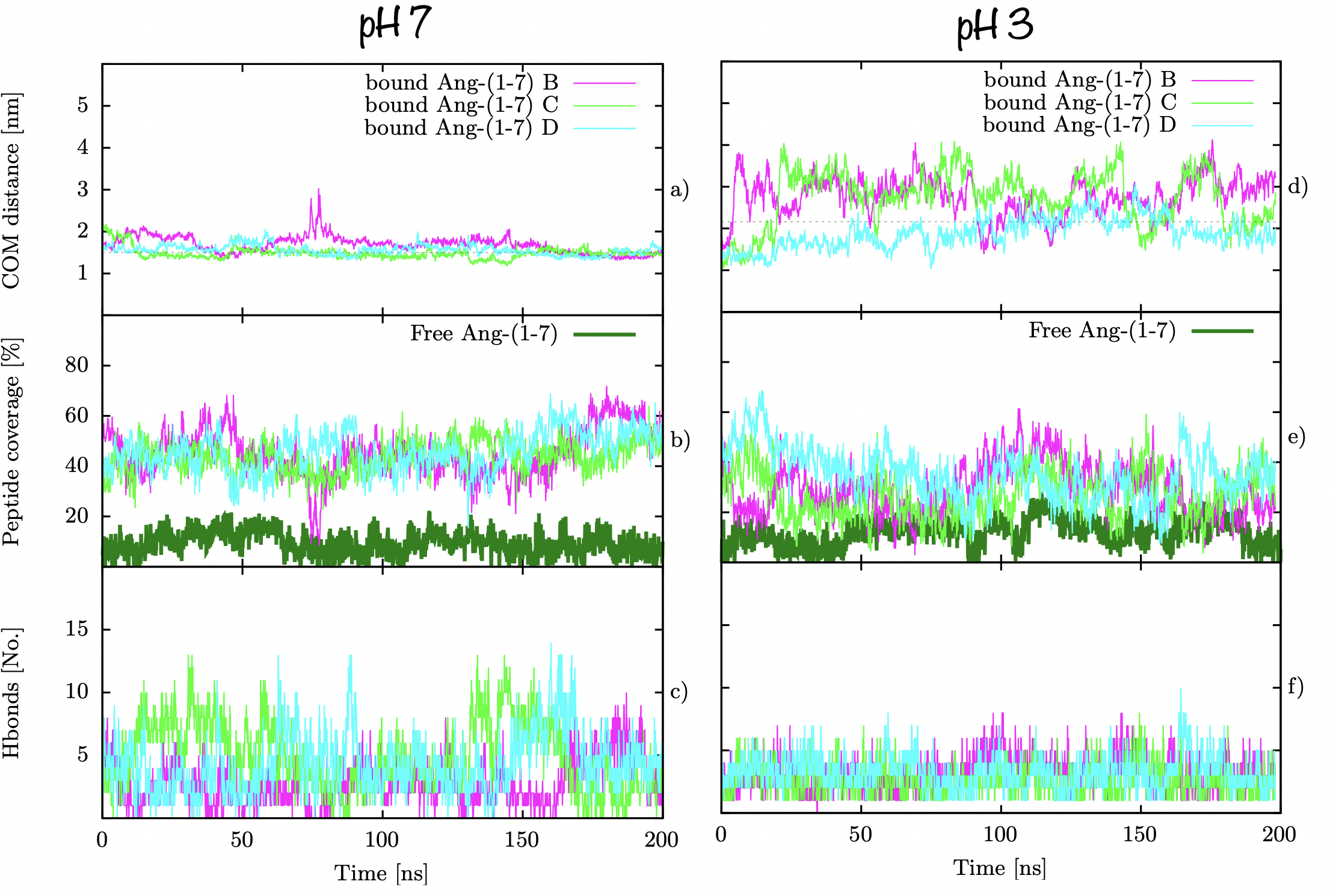

### contac_map.pdf

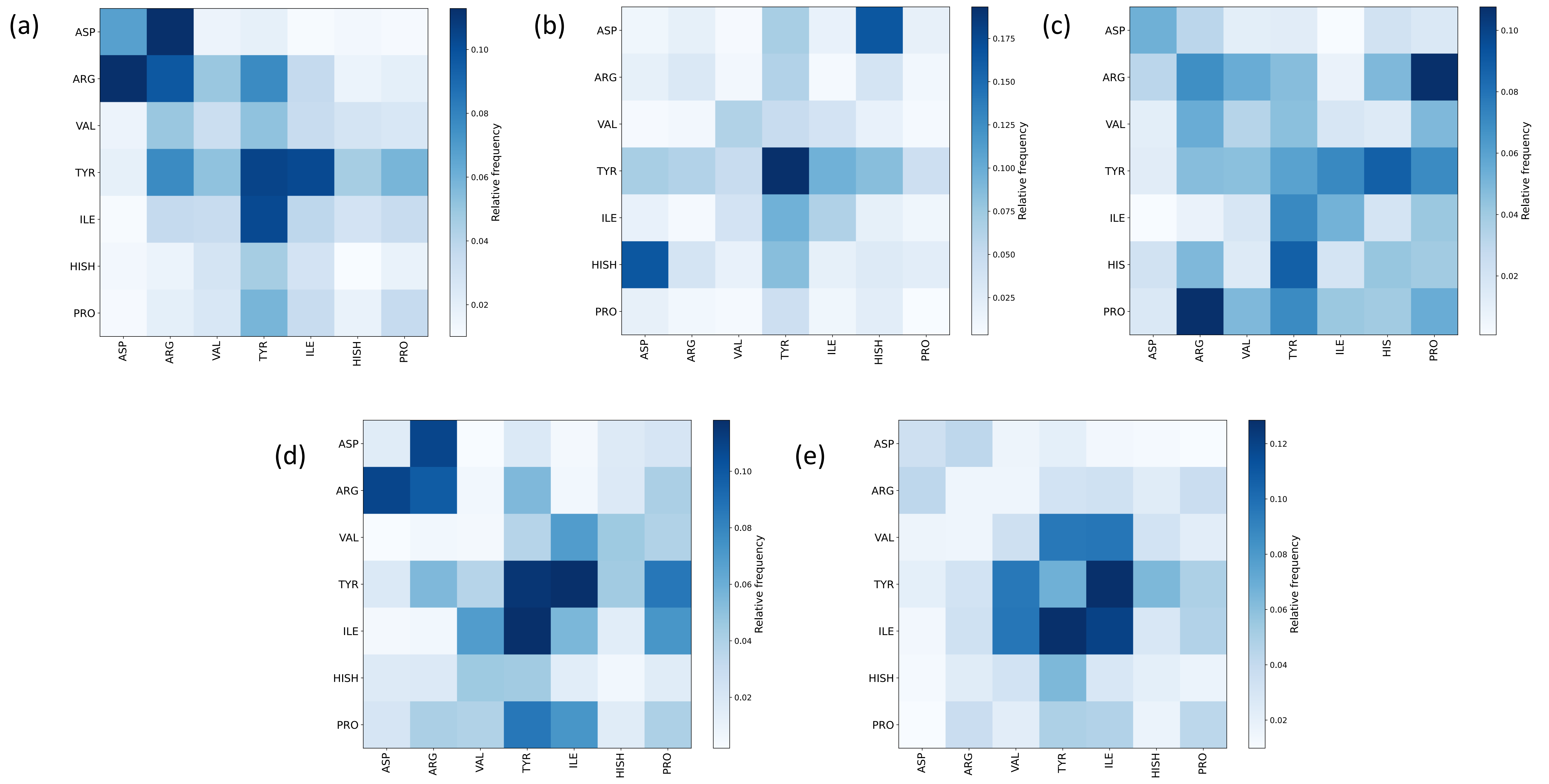

### contributions2.pdf

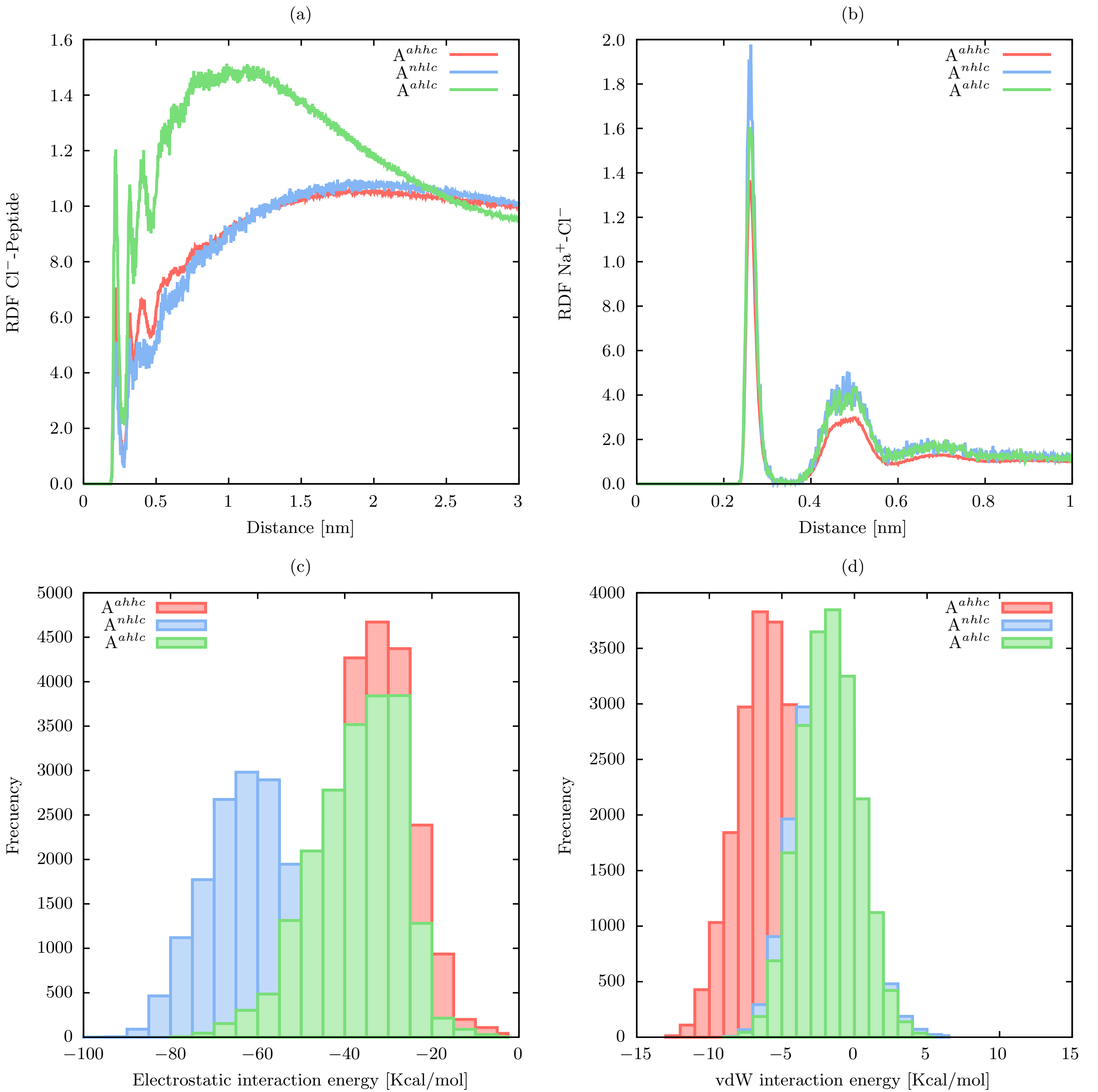

### contributions.pdf

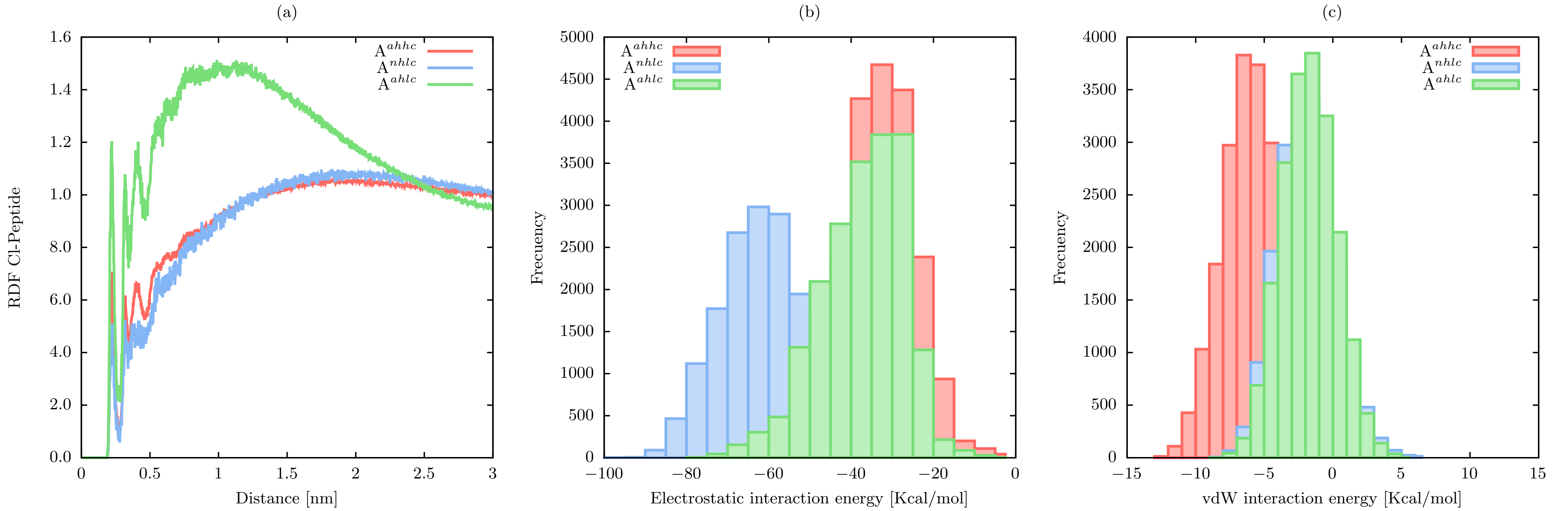

### coordinations_pnhcom.jpg

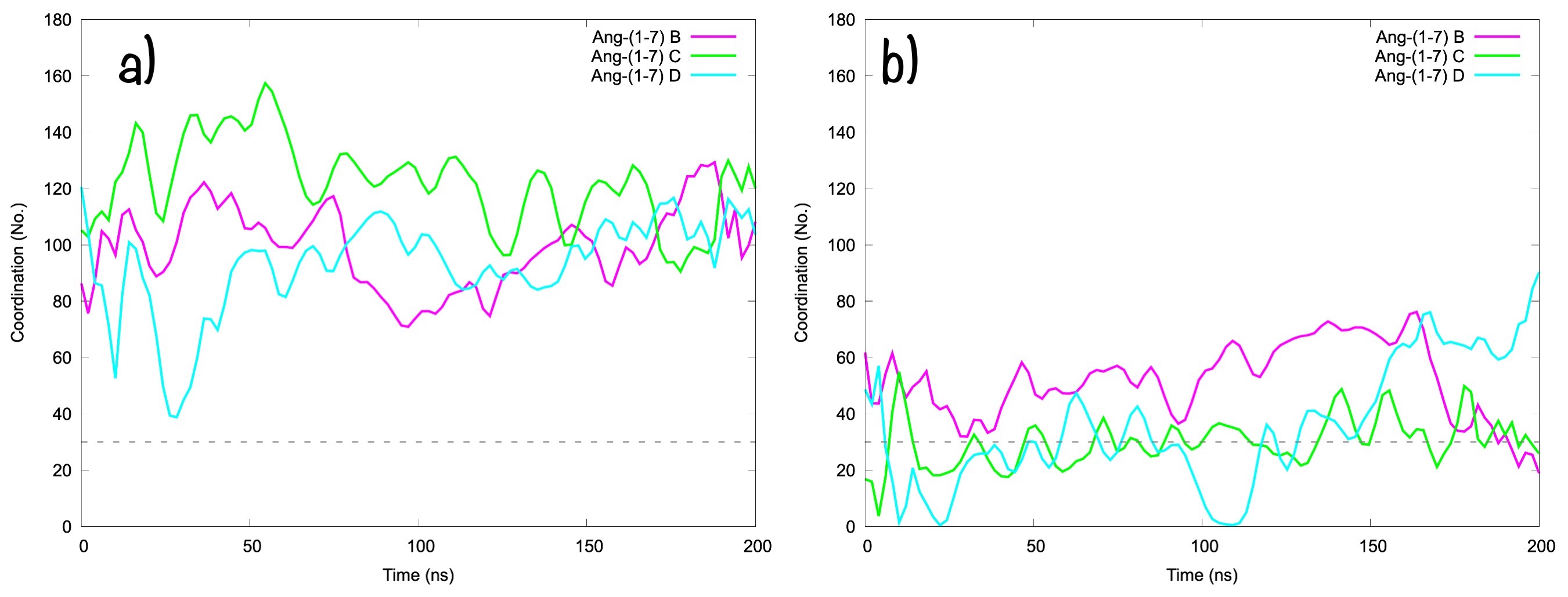

### coordinations_pohcom.jpg

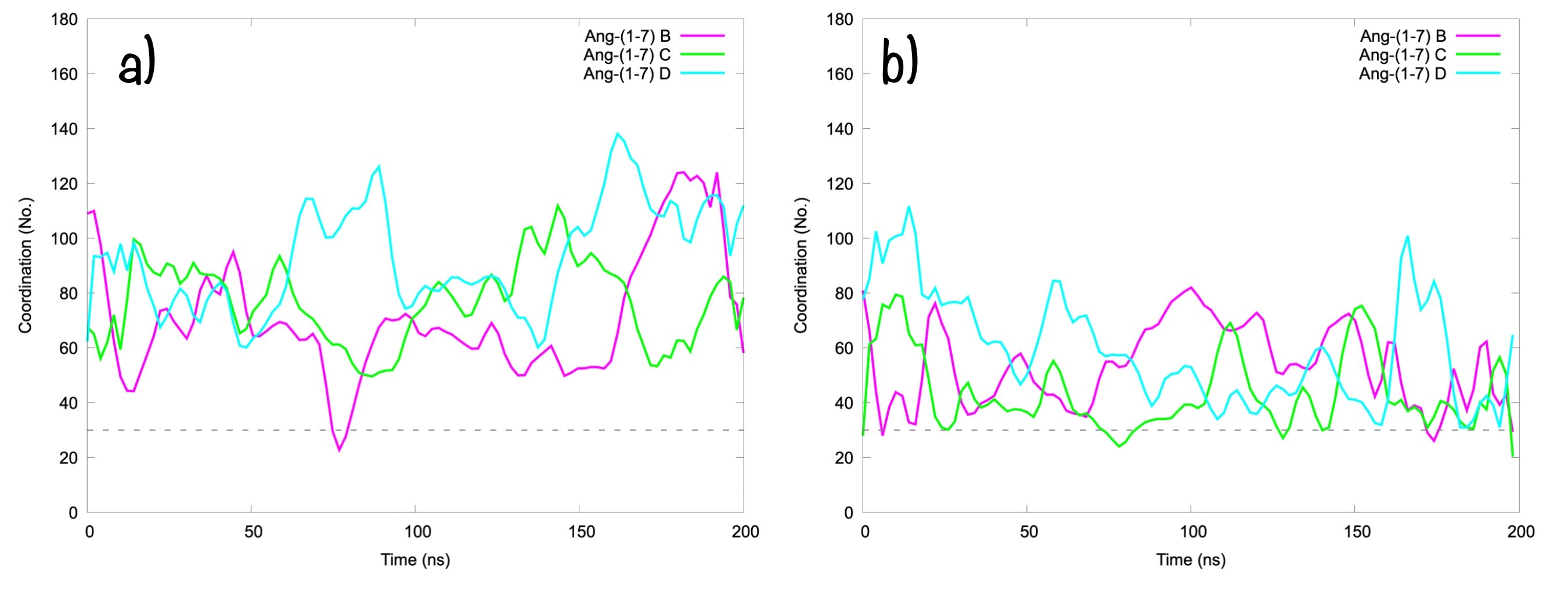

### dock.png

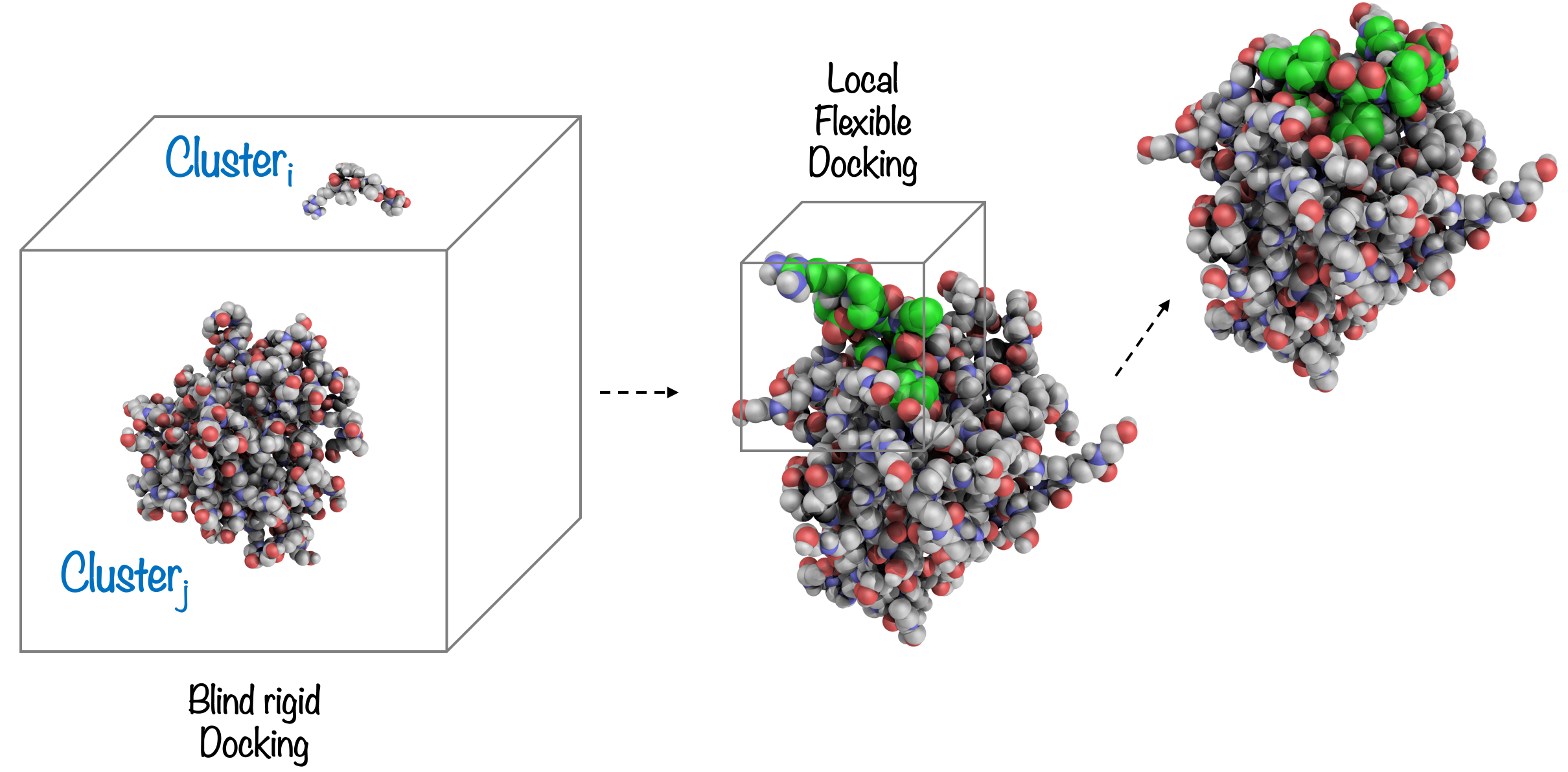

### fes-ang-ph7.png

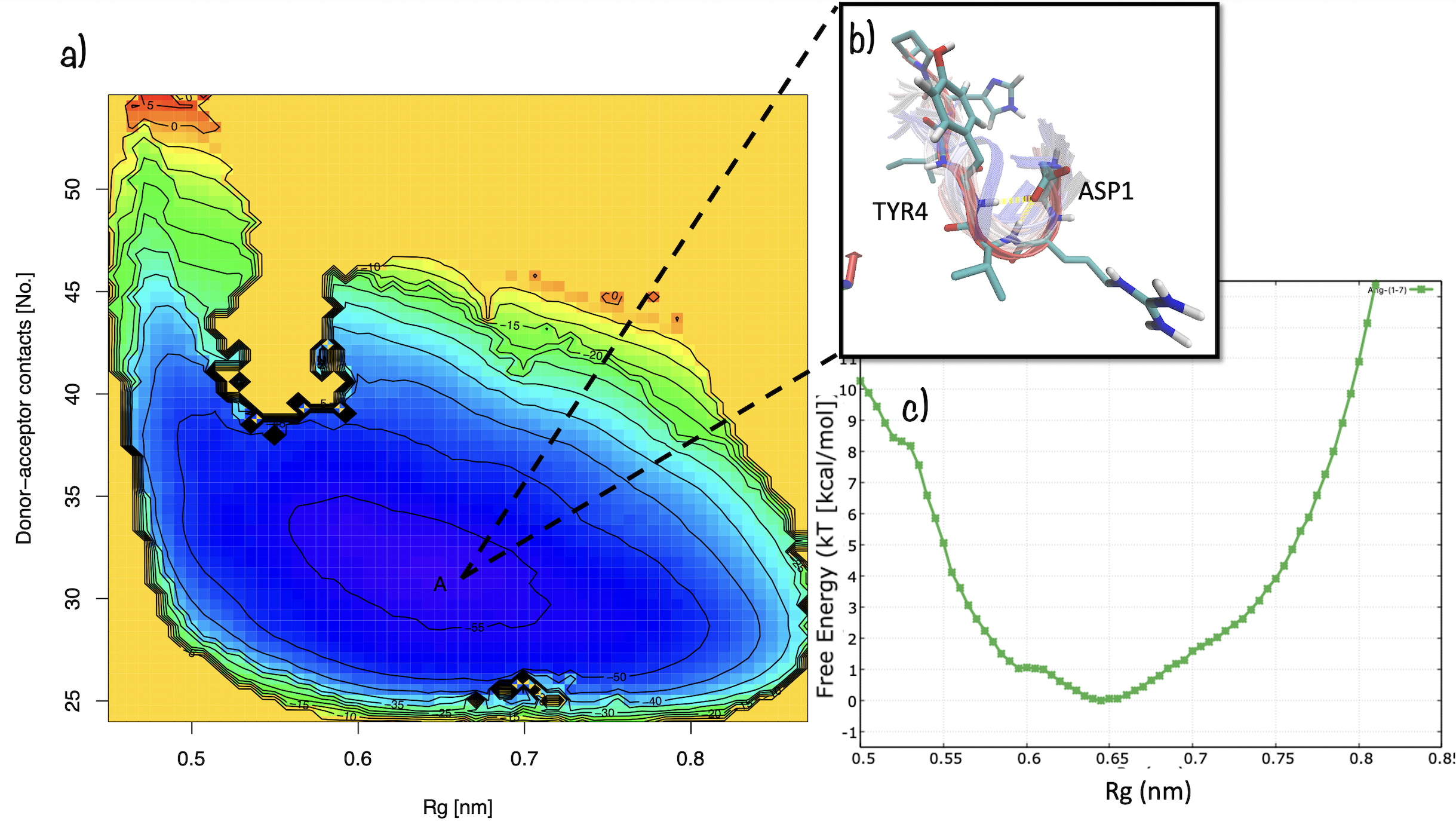

### gyrate_all_1.pdf

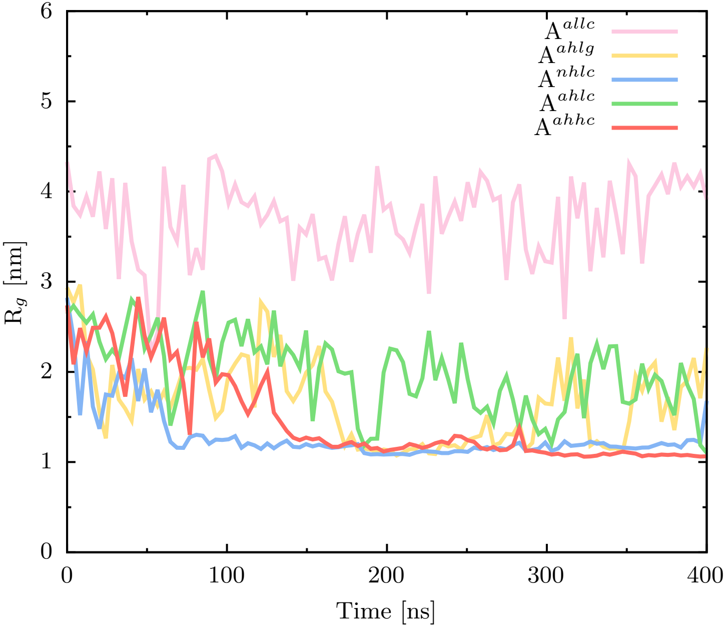

### gyration.pdf

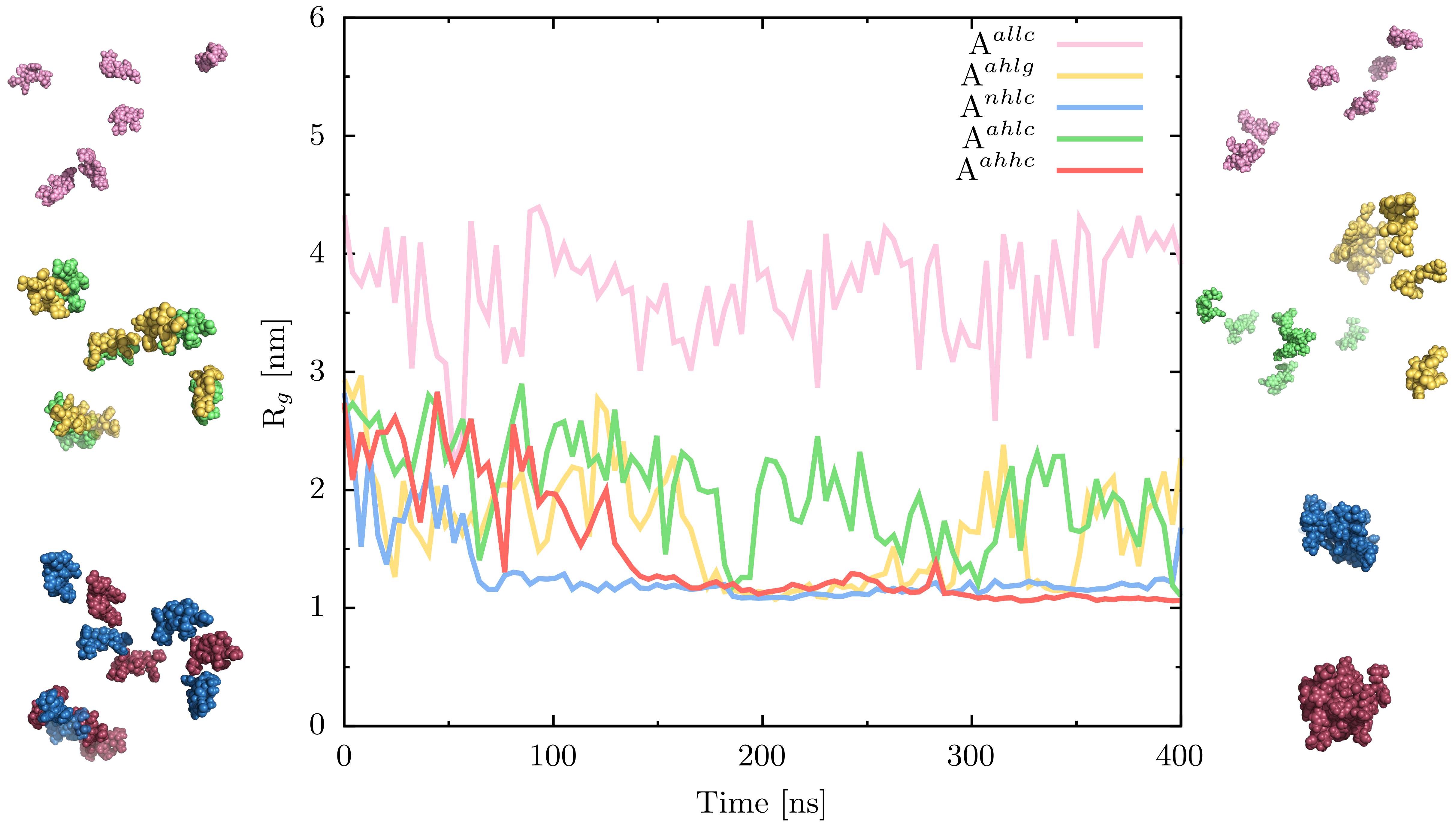

### md_properties_ph3-7_200ns-1.pdf

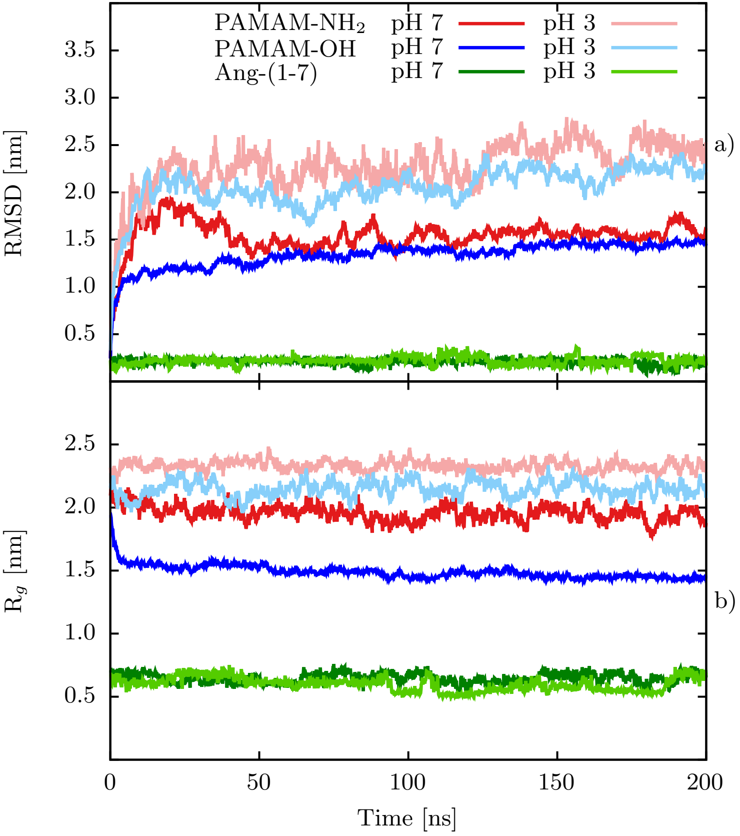

### md_properties_ph7_200ns-1.pdf

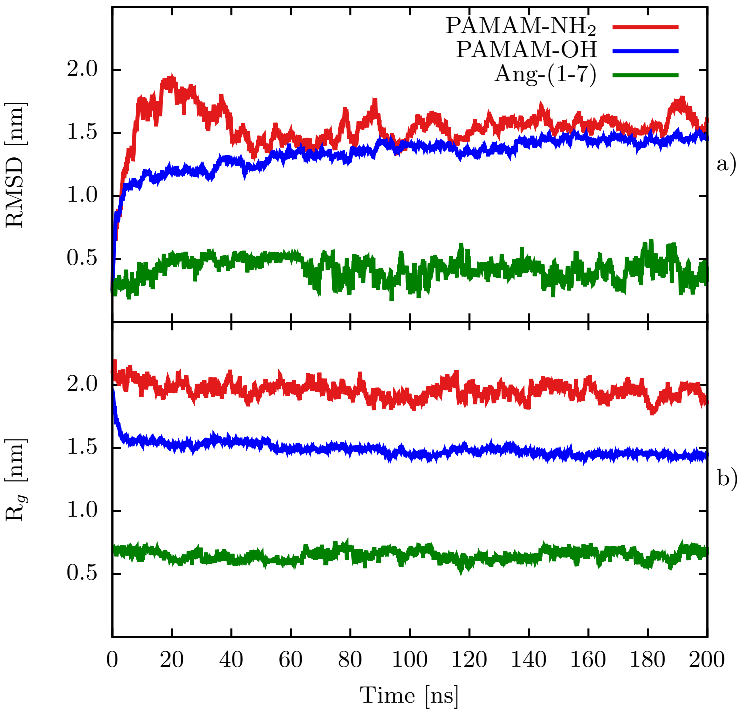

### metady-ph7-clusters.png

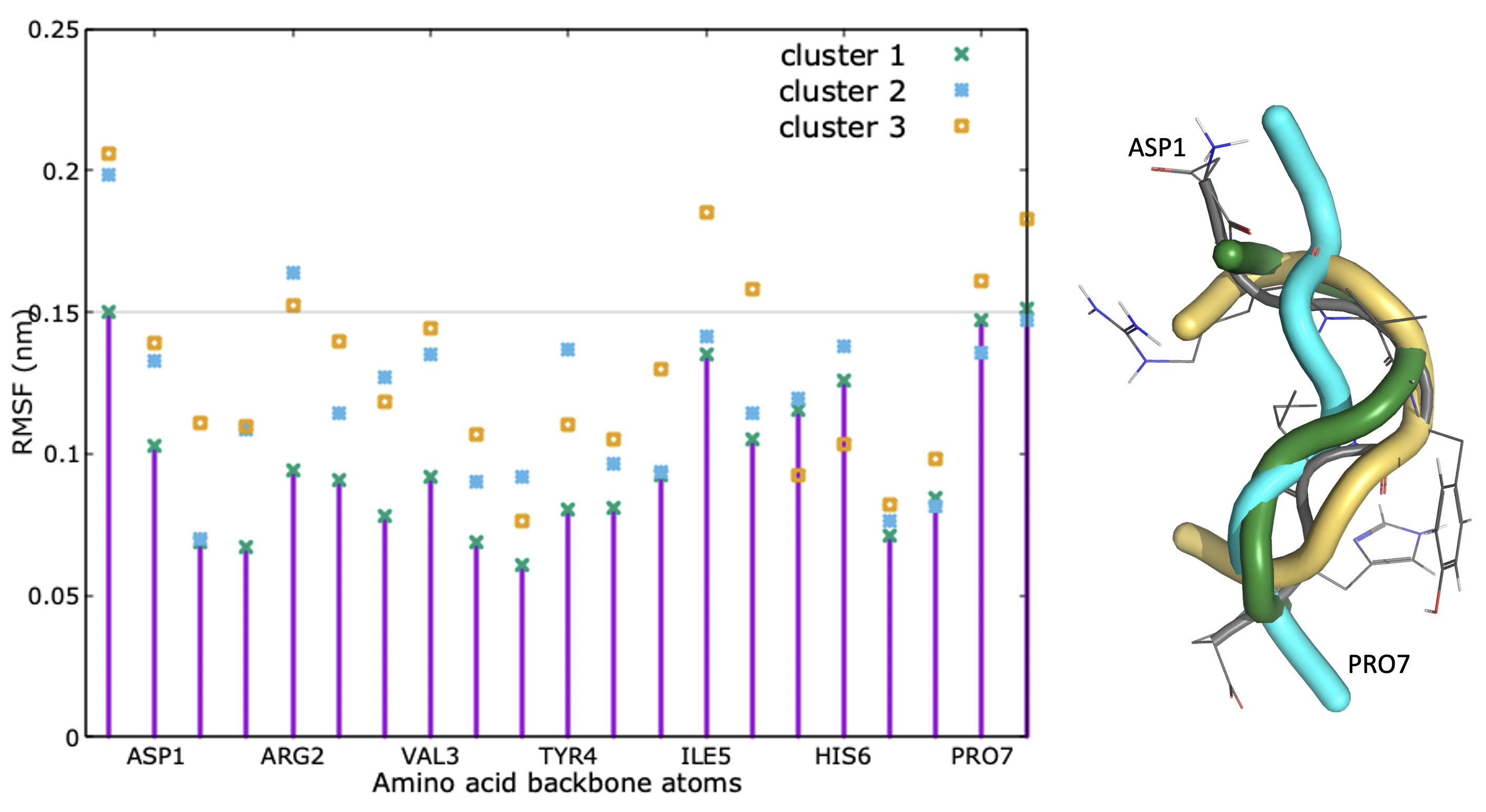

### multiligand-complexes.png

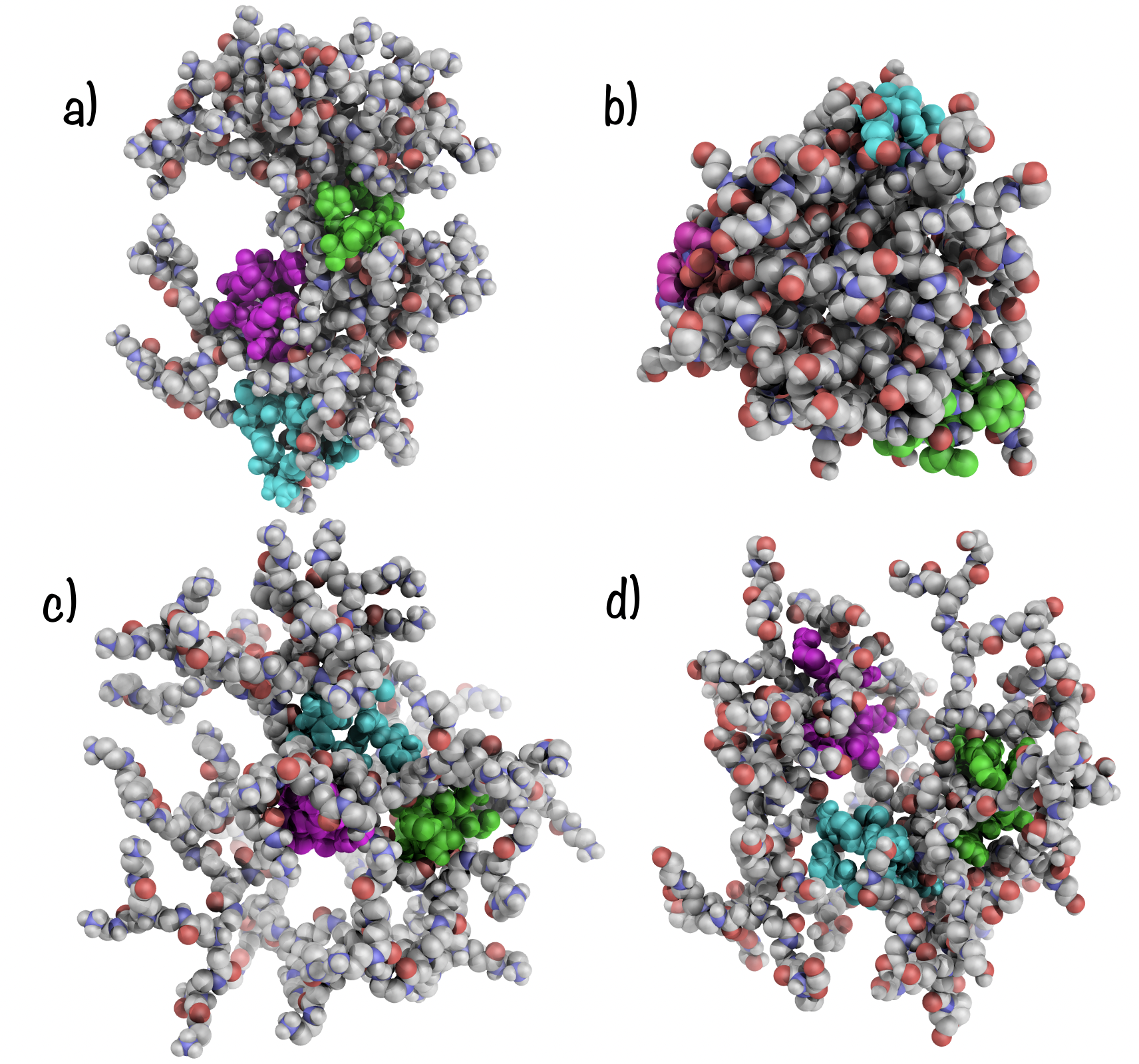

### ocu-pnh-struc.png

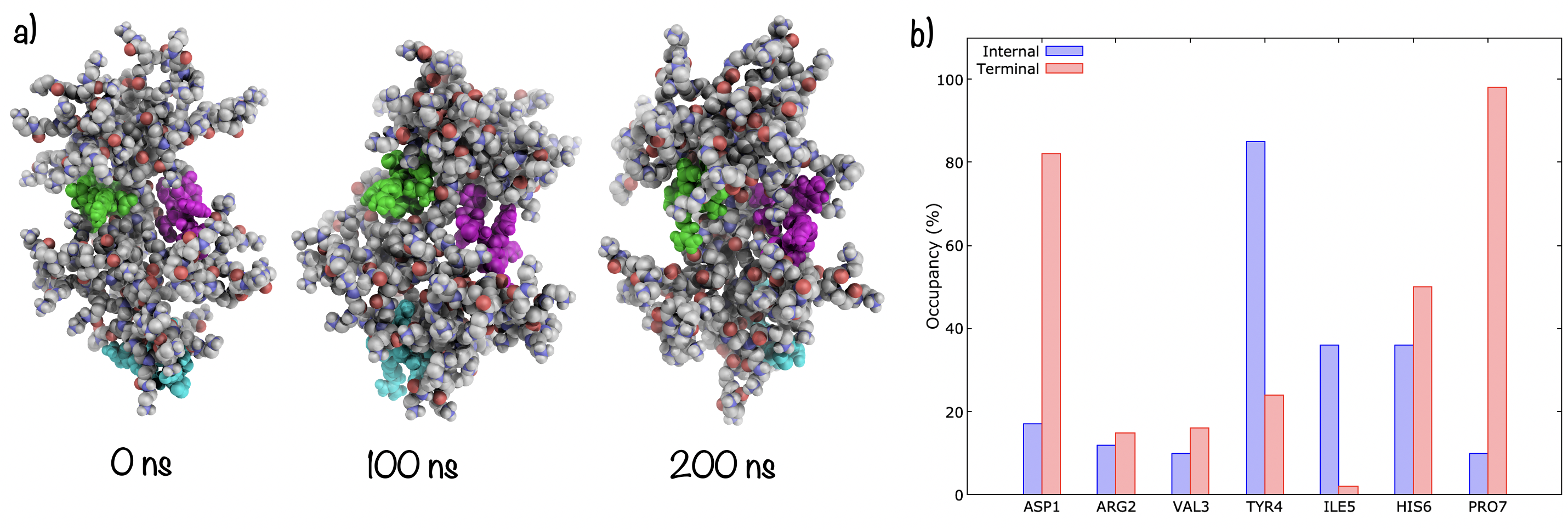

### ocu-poh-2.png

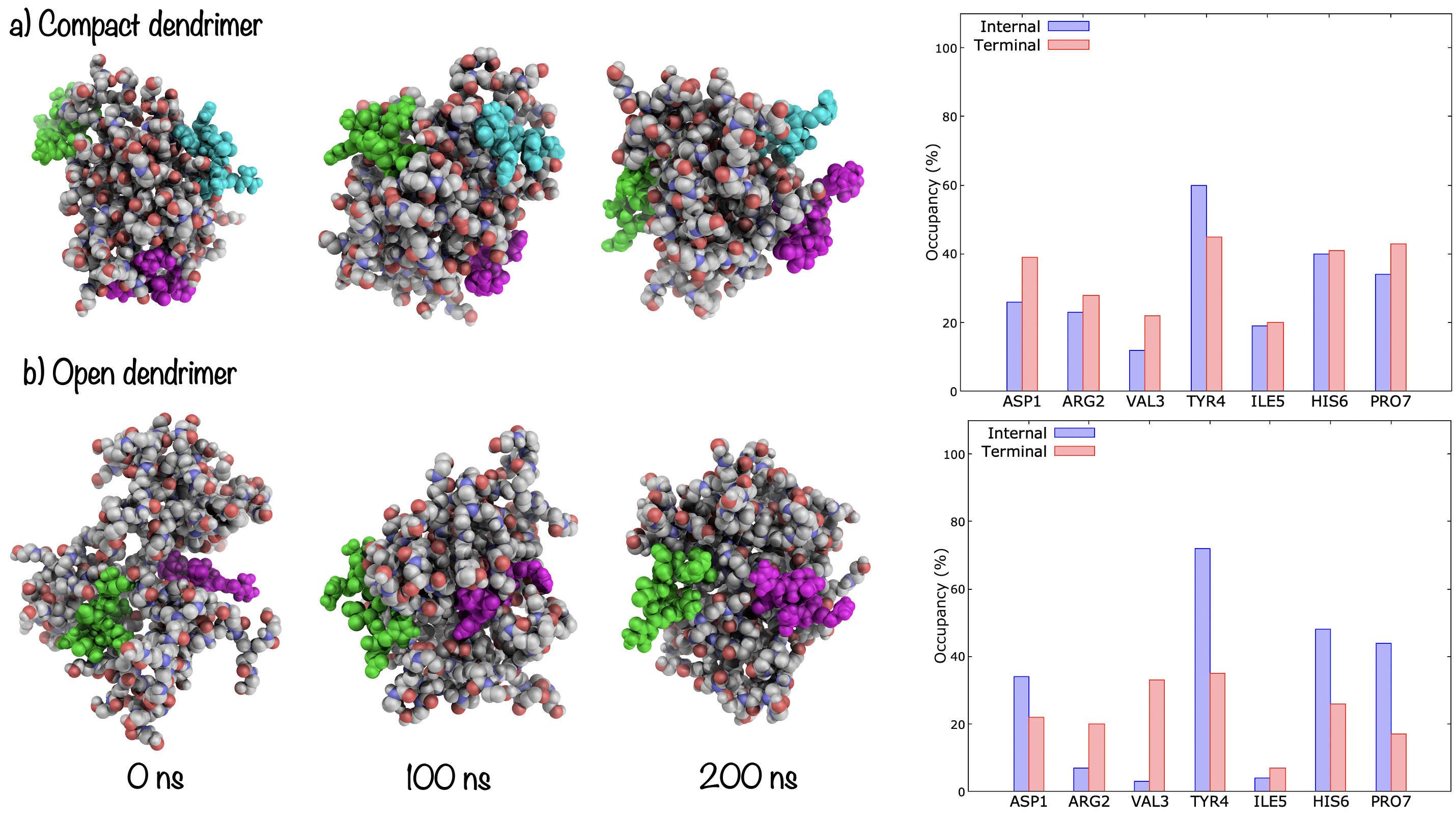

### pamamnhcom-hist.png

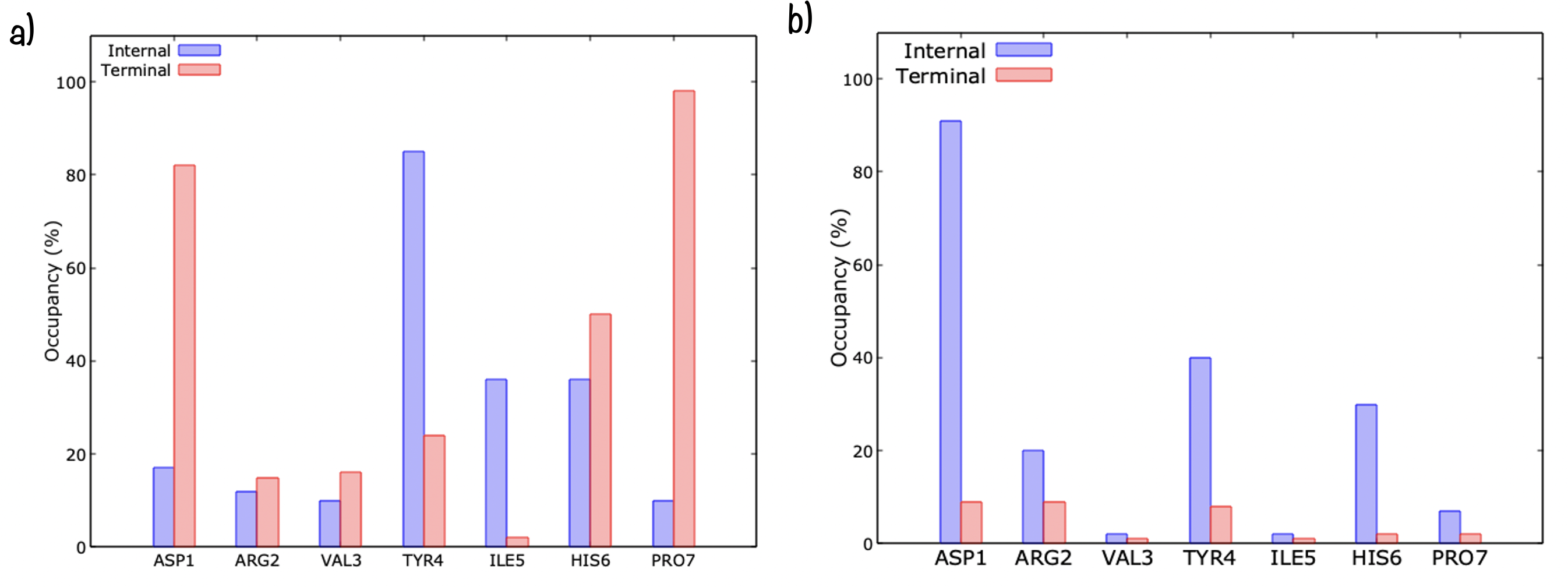

### pamamohcom-hist.png

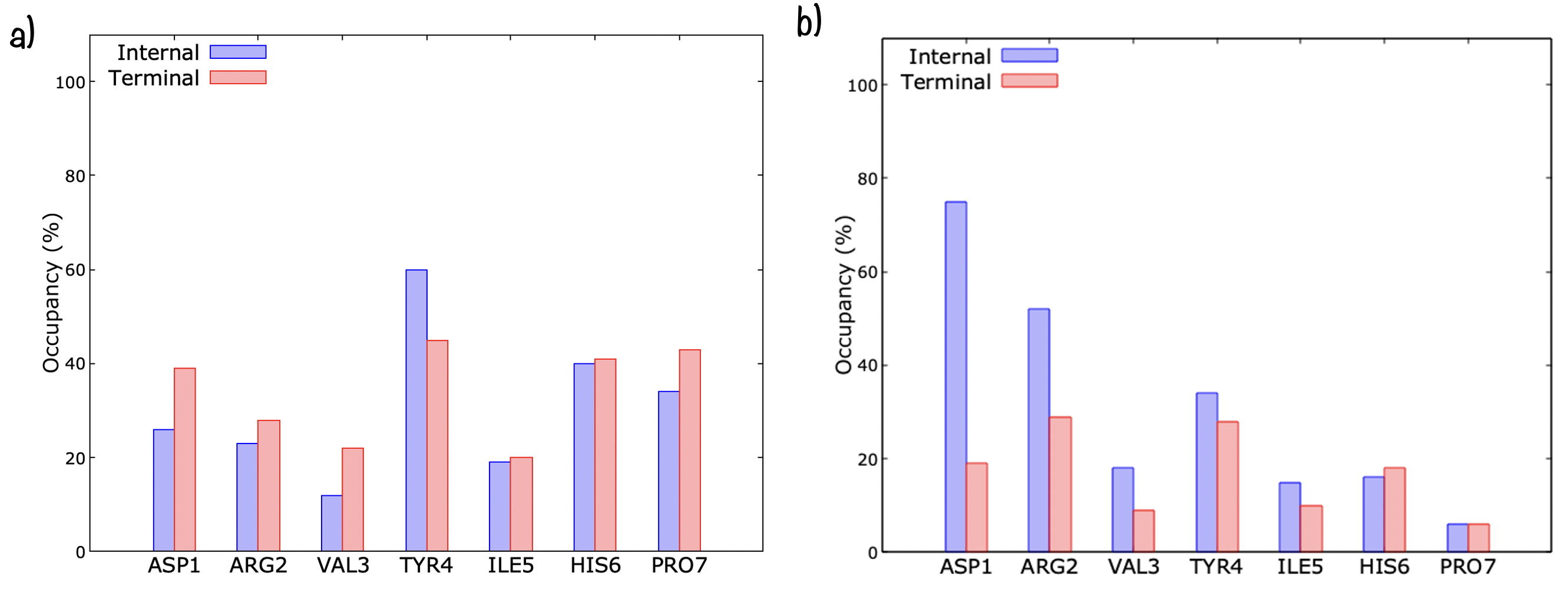

### pathway.png

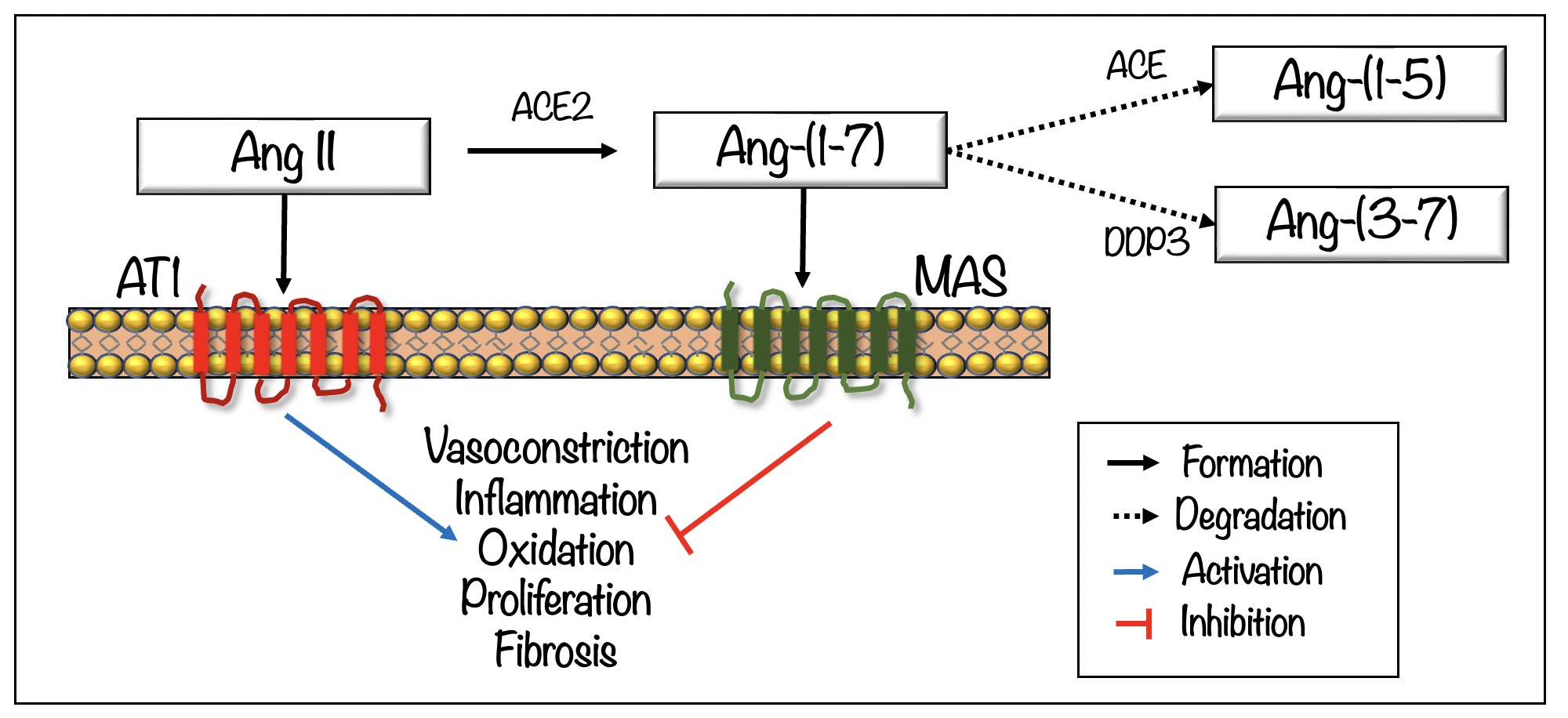

### pnh3com.png

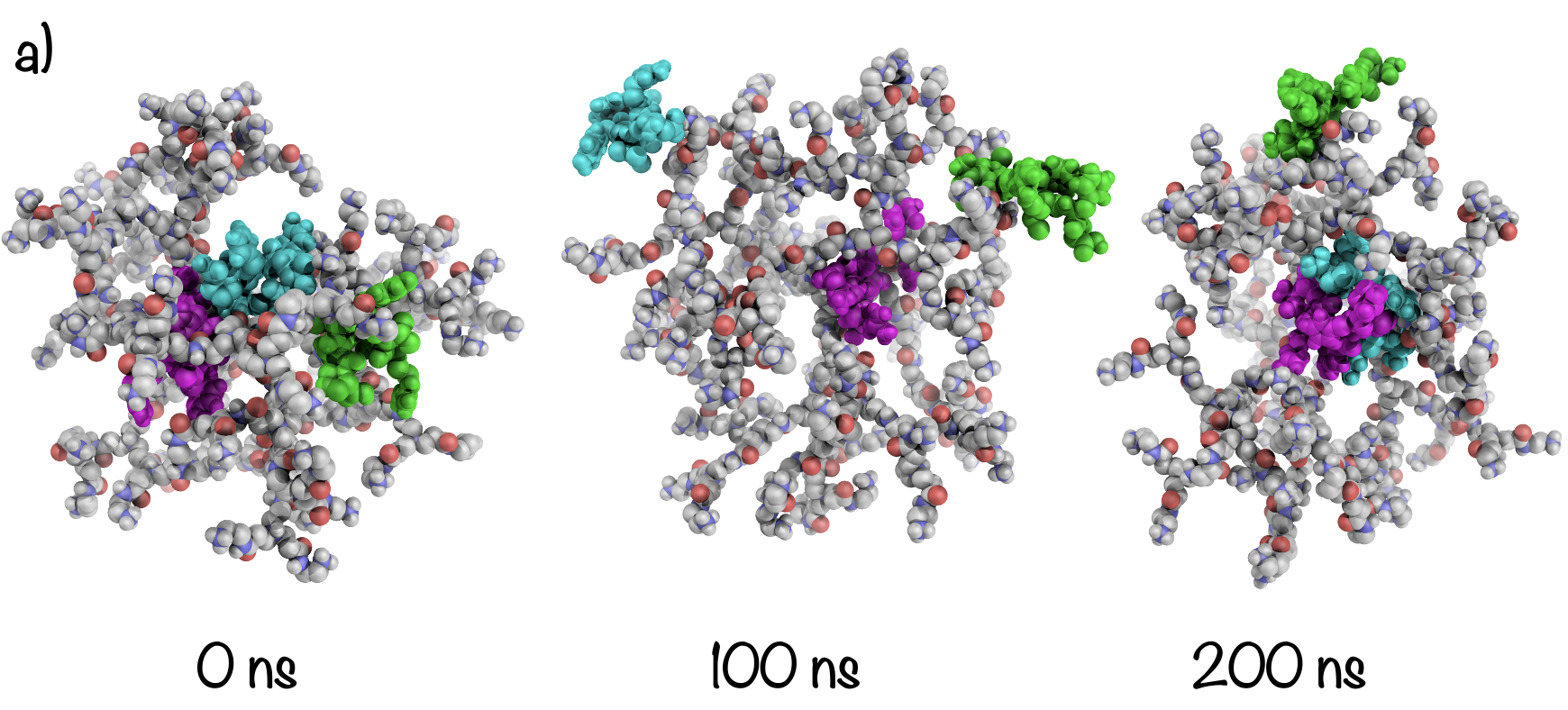

### proteolitic.png

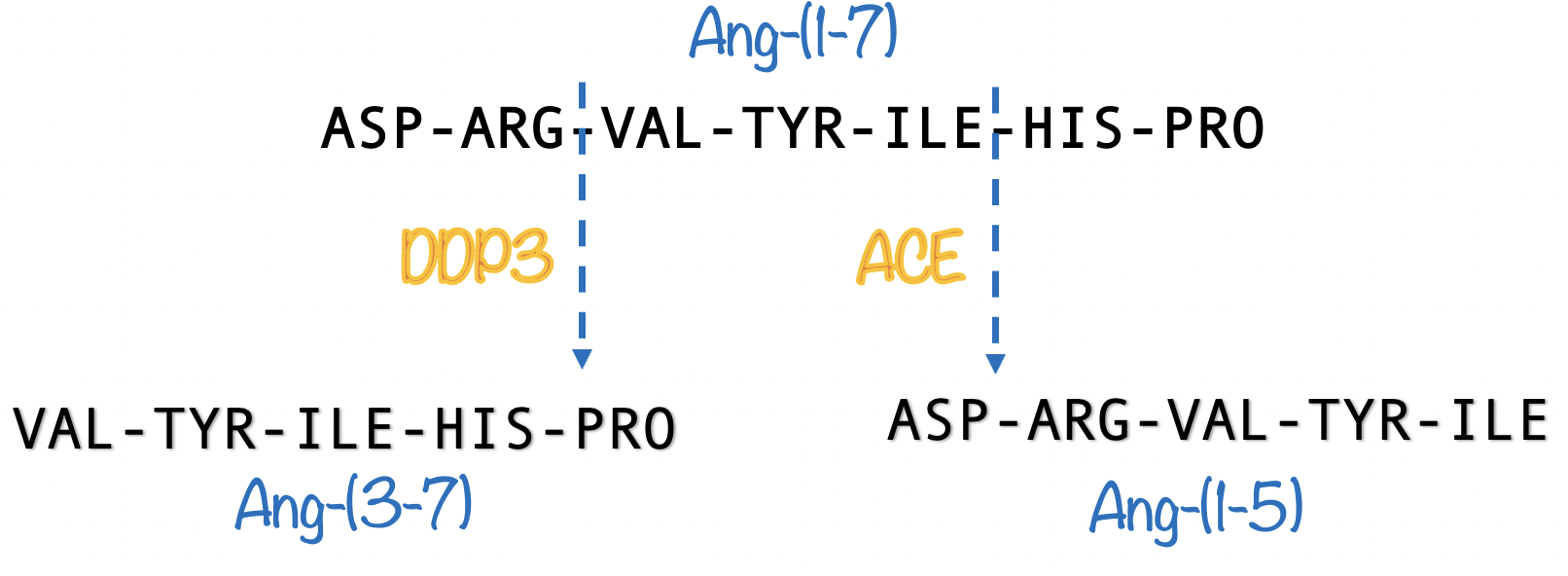

### rdf-1.png

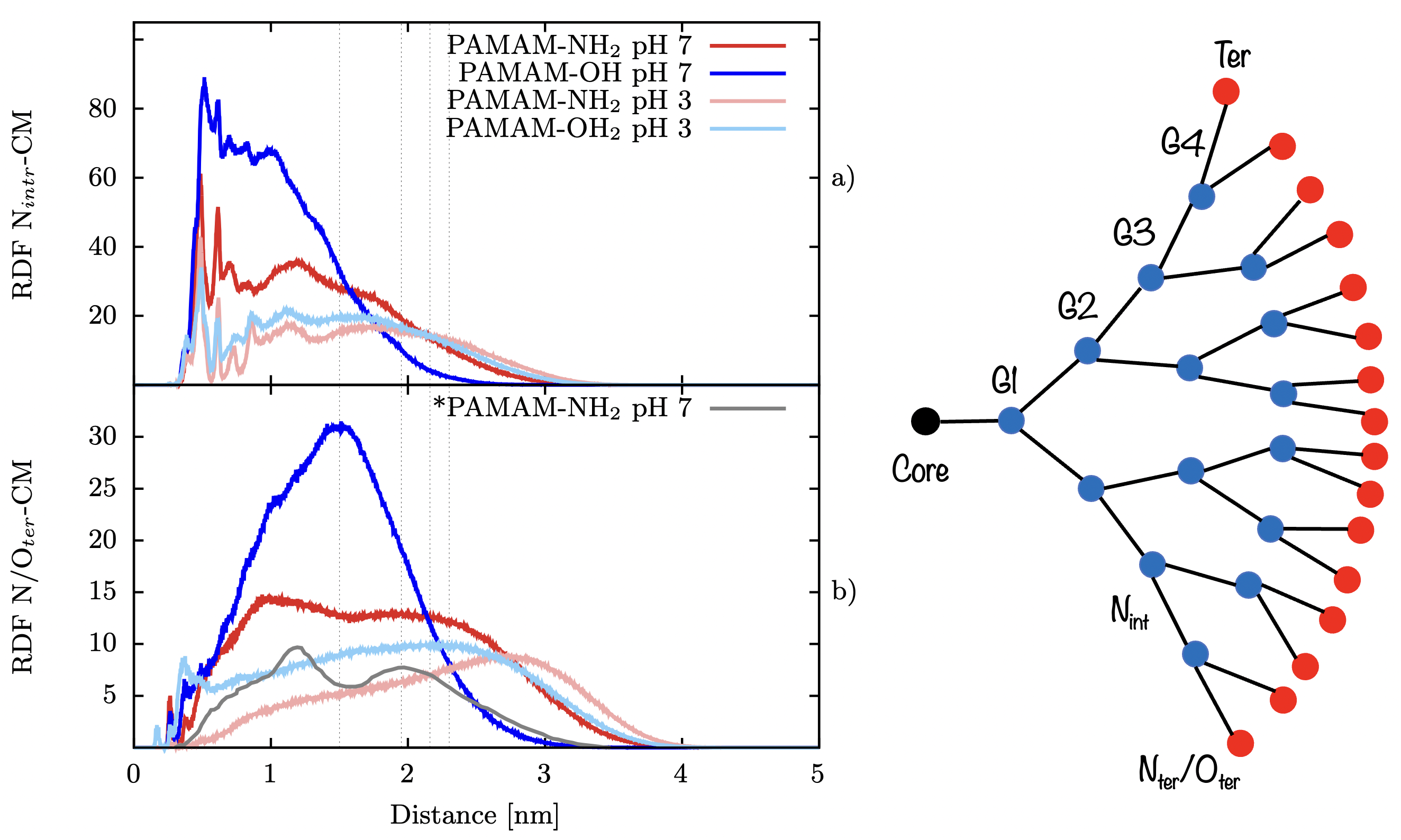

### rdf-2.png

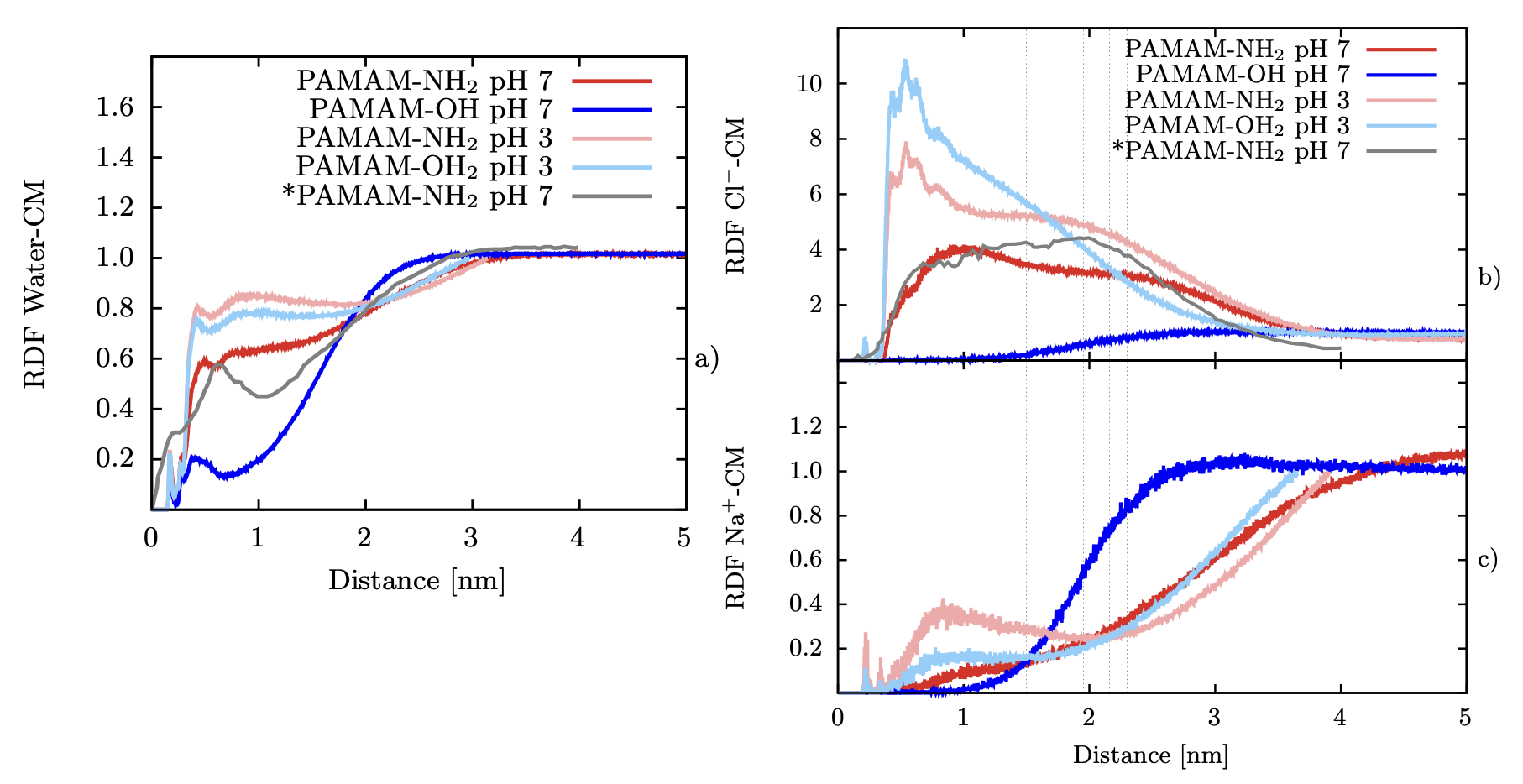

### sphericity-ph3-ph7.png

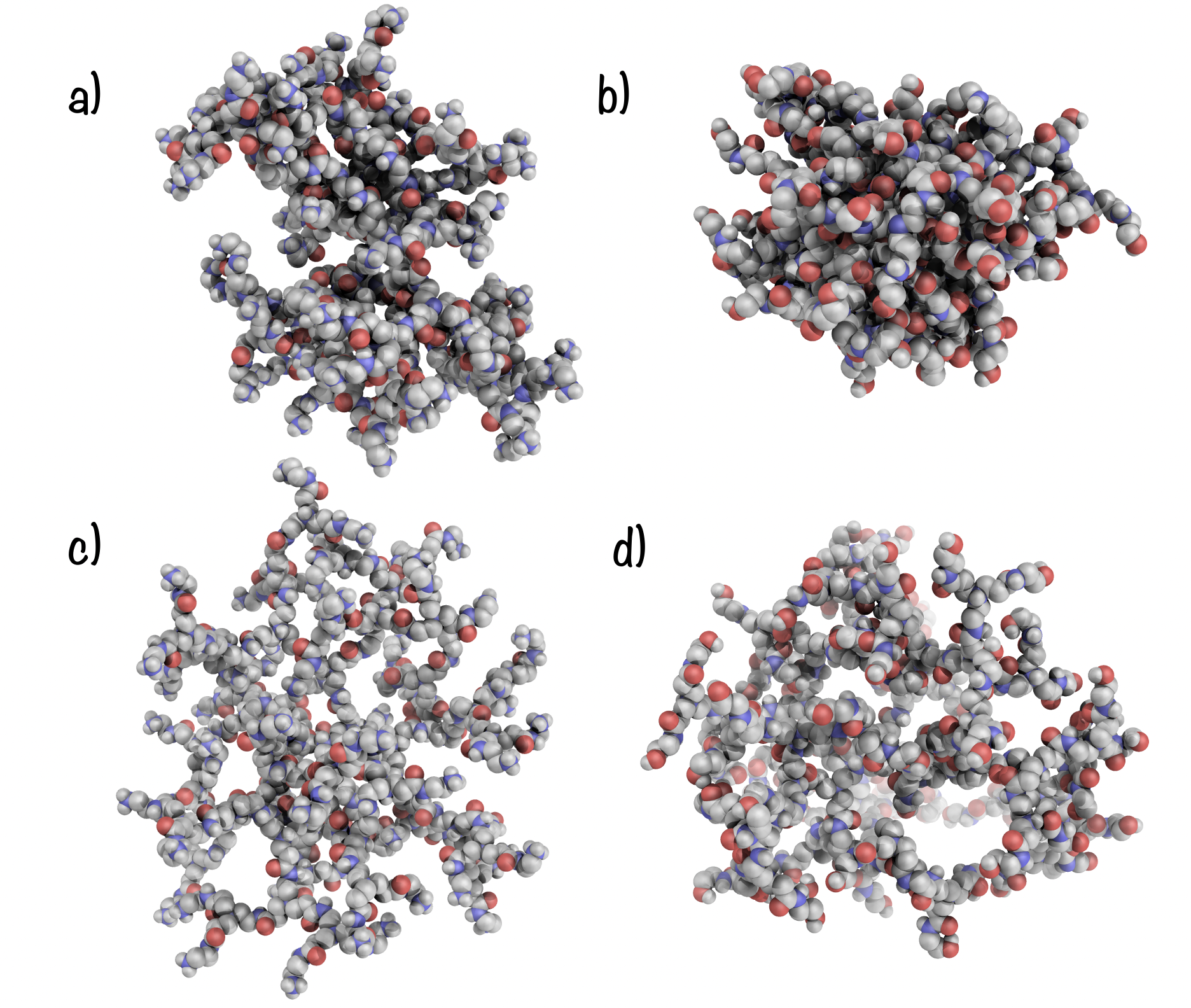

### superimposed_ang.png

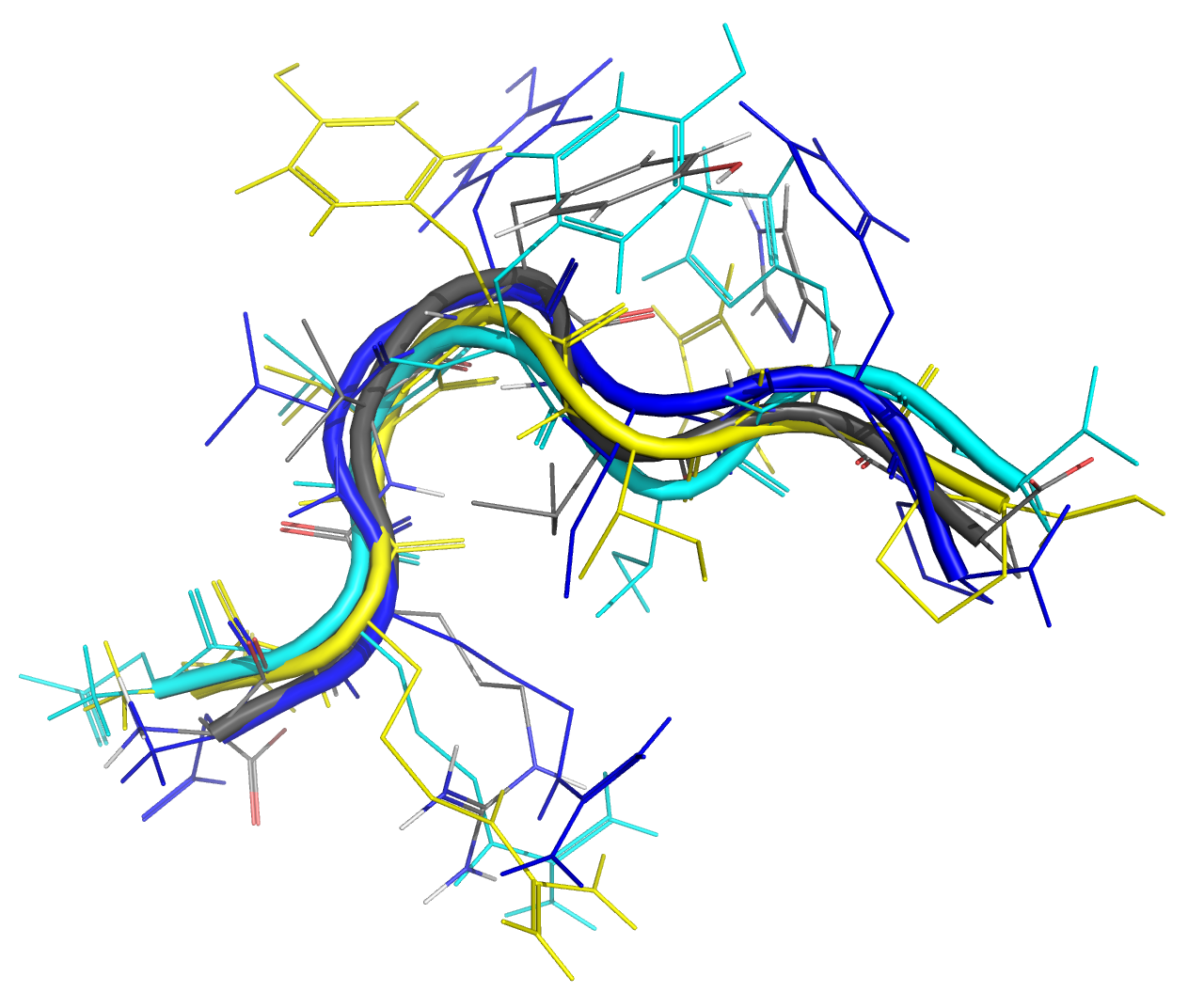

### ter-wat-den.png

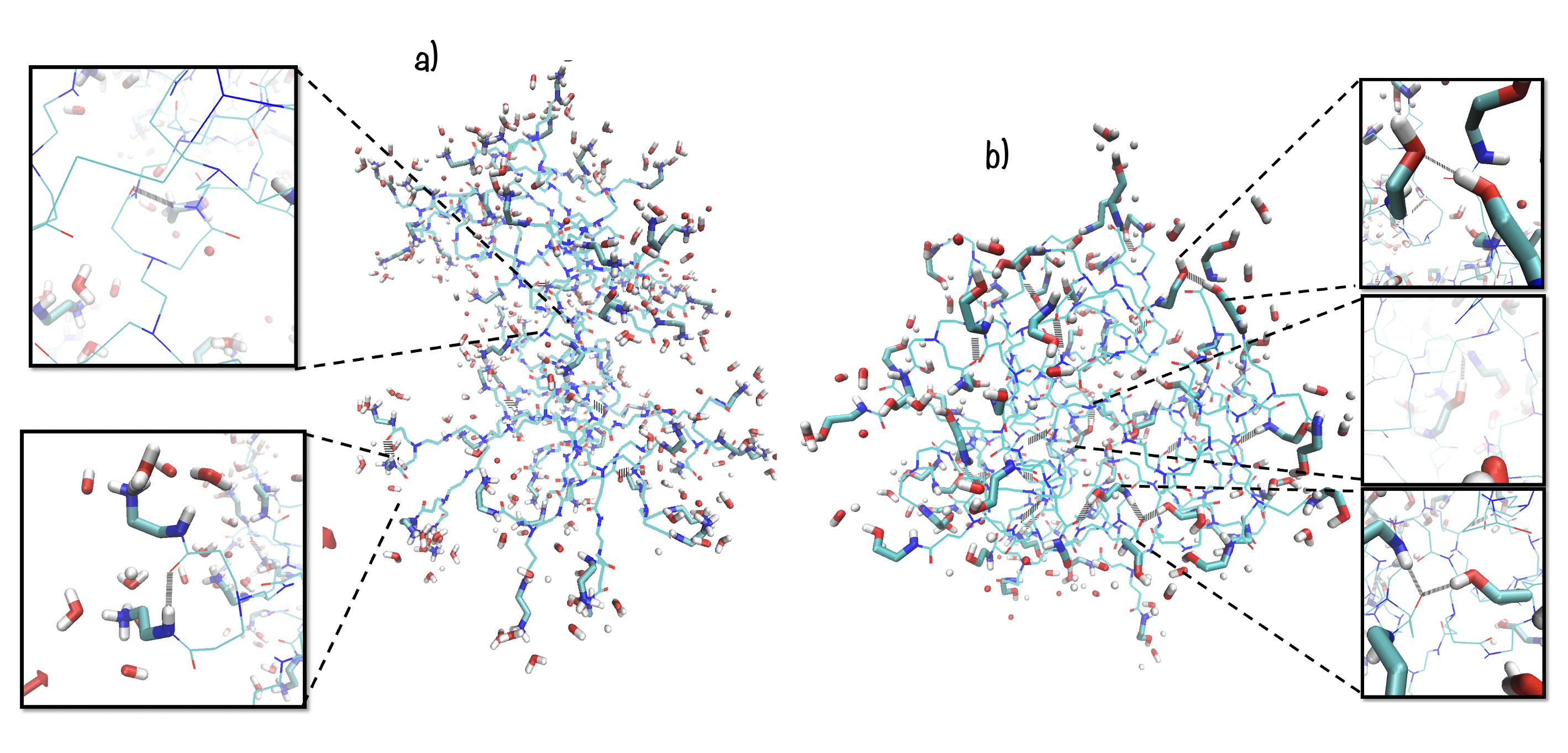

### timeline-pnhcom.png

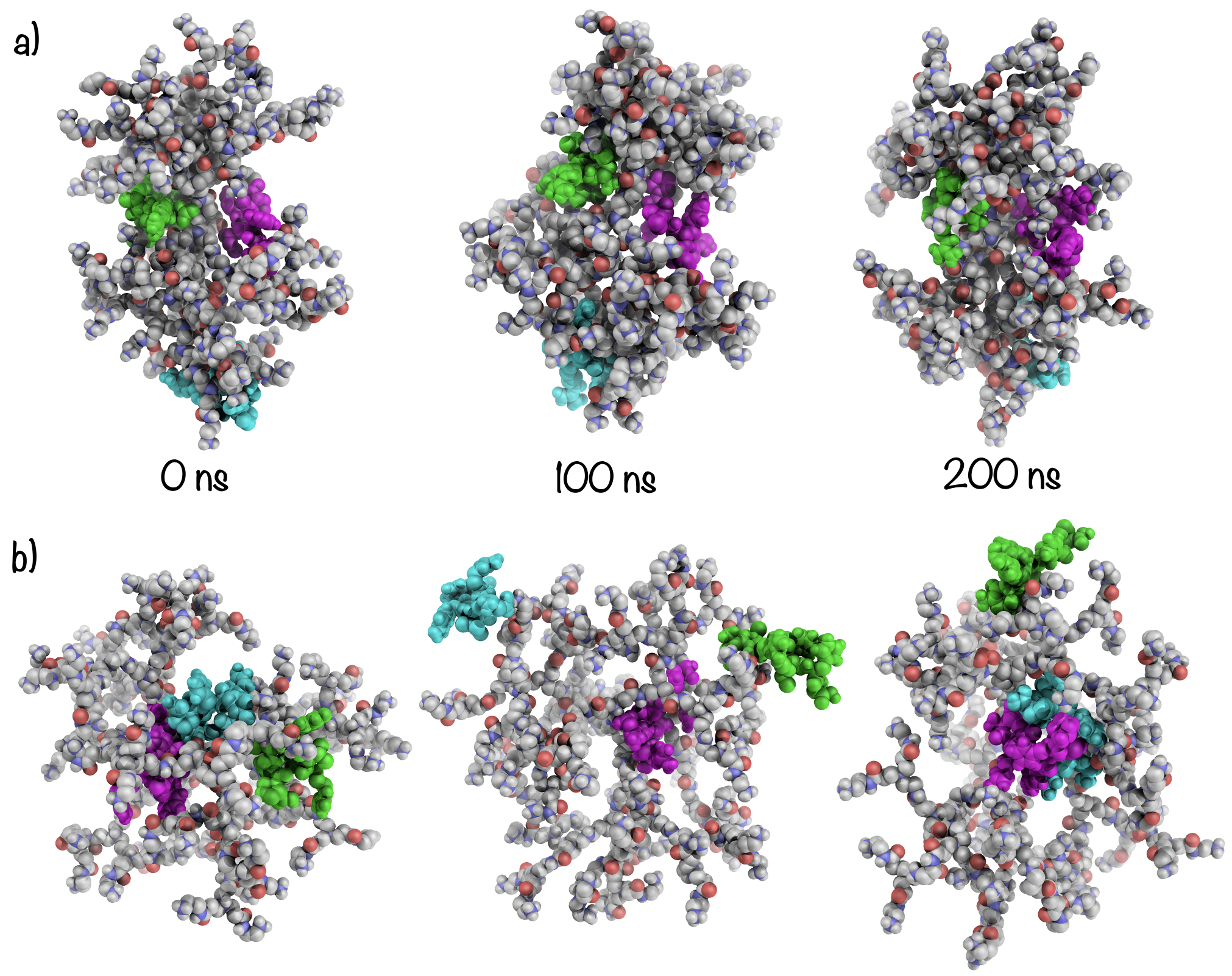

### timeline-pohcom.png

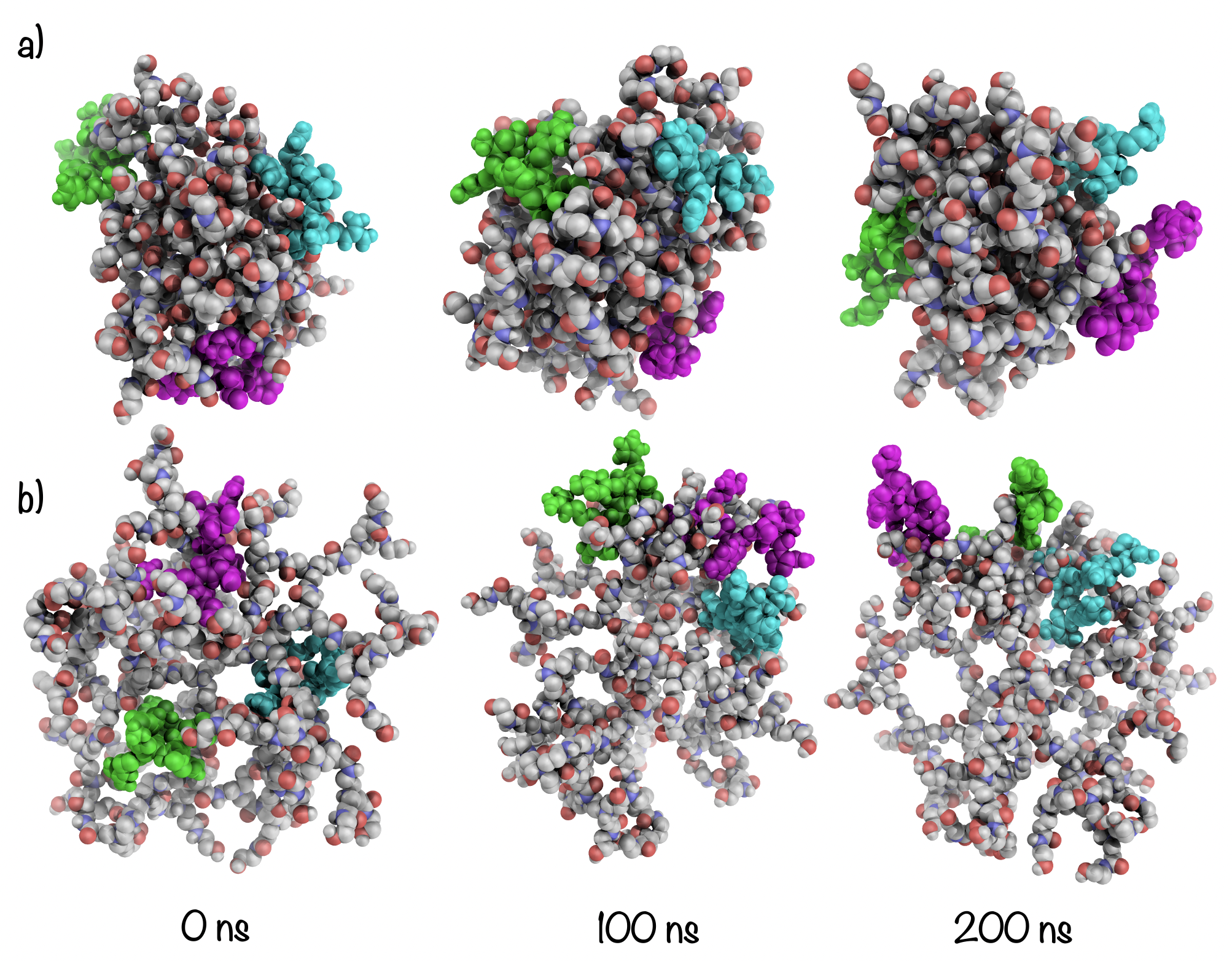
